# The *Zea mays* PeptideAtlas – a new maize community resource

**DOI:** 10.1101/2023.12.21.572651

**Authors:** Klaas J. van Wijk, Tami Leppert, Zhi Sun, Isabell Guzchenko, Erica Debley, Georgia Sauermann, Pratyush Routray, Luis Mendoza, Qi Sun, Eric W. Deutsch

**Author notes:** correspondence: Klaas J. van Wijk,; Eric W. Deutsch.

## Abstract

We developed the Maize PeptideAtlas resource (www.peptideatlas.org/builds/maize) to help solve questions about the maize proteome. Publicly available raw tandem mass spectrometry (MS/MS) data for maize were collected from ProteomeXchange and reanalyzed through a uniform processing and metadata annotation pipeline. These data are from a wide range of genetic backgrounds, including the inbred lines B73 and W22, many hybrids and their respective parents. Samples were collected from field trials, controlled environmental conditions, a range of (a)biotic conditions and different tissues, cell types and subcellular fractions. The protein search space included different maize genome annotations for the B73 inbred line from MaizeGDB, UniProtKB, NCBI RefSeq and for the W22 inbred line. 445 million MS/MS spectra were searched, of which 120 million were matched to 0.37 million distinct peptides. Peptides were matched to 66.2% of the proteins (one isoform per protein coding gene) in the most recent B73 nuclear genome annotation (v5). Furthermore, most conserved plastid- and mitochondrial-encoded proteins (NCBI RefSeq annotations) were identified. Peptides and proteins identified in the other searched B73 genome annotations will aid to improve maize genome annotation. We also illustrate high confidence detection of unique W22 proteins. N-terminal acetylation, phosphorylation, ubiquitination, and three lysine acylations (K-acetyl, K-malonyl, K-hydroxyisobutyryl) were identified and can be inspected through a PTM viewer in PeptideAtlas. All matched MS/MS-derived peptide data are linked to spectral, technical and biological metadata. This new PeptideAtlas is integrated with community resources including MaizeGDB at https://www.maizegdb.org/ and a peptide track in JBrowse.

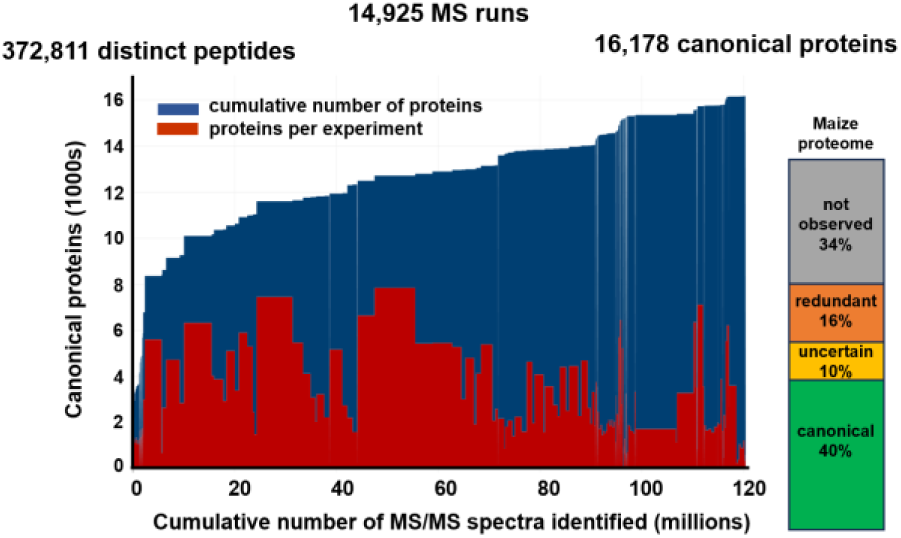

## INTRODUCTION

Maize is a crop of major agricultural importance and also serves as an important model system for the study of grasses and C4 photosynthesis ^1^. Different cultivars and hybrids are grown for agriculture but the B73 inbred was initially chosen for maize genome sequencing and it has become the reference maize line with the most genomic resources for the research community ^2–5^. After the first maize B73 genome assembly in 2009 ^2^, several subsequent B73 assemblies followed in rapid order with RefGen_v5 being the most recent ^6, 7^. In the last few years, additional maize genomes have been sequenced and assembled for more than 25 inbred lines, including W22 ^8^, RP125 ^9^, A188 ^10^, and 25 maize nested association mapping (NAM) founder inbred lines^7^.

These different annotated maize genomes each predict over 38000 protein coding genes across the 10 maize nuclear chromosomes, and these genes are represented by a larger number of transcripts. Many aspects of the cellular proteome cannot be predicted but must be experimentally determined at the protein level, such as cell-type specific and subcellular protein accumulation, protein post-translational modifications (PTMs), half-life, and protein interactions. Furthermore, even the best annotated genomes cannot easily predict which mRNA splice forms result in proteins, and MS workflows face technical challenges to identify all possible peptide covering splice junctions, see *e.g.* ^11, 12^. Therefore, the impact of alternative splicing for the cellular proteome in eukaryotes is still under debate ^13–15^. In addition, plant genomes may also contain an unknown number of various types of small Open Reading Frames (sORFs) encoding for small proteins or peptides, that could be detected by MS ^16–18^. The use of proteomics data for plant genome annotation is summarized under the term ‘proteogenomics’ ^19^, and has been applied for the genomes of Arabidopsis (*Arabidopsis thaliana*) ^20, 21^, rice (*Oryza sativa*) ^22, 23^, maize ^24^, grape (*Vitis vinifera*) ^25^, and sweet potato (*Ipomoea batatas*) ^26^ and the perennial fruit cherry ^27^.

Recently, we published two studies that describes a new community proteomics resource for *Arabidopsis thaliana* entitled Arabidopsis PeptideAtlas, http://www.peptideatlas.org/builds/arabidopsis/. ^17, 28^. The purpose of this new resource is to help solve central questions about the Arabidopsis proteome, such as the significance of protein splice forms, post-translational modifications (PTMs), and obtain reliable information about specific proteins. PeptideAtlas is based on published mass spectrometry (MS) datasets collected through ProteomeXchange (http://www.proteomexchange.org/) and reanalyzed with the Trans-Proteomic Pipeline (TPP) ^29–31^ and our metadata annotation pipeline. Arabidopsis PeptideAtlas is integrated with community resources including TAIR, JBrowse, PPDB and UniProtKB.

In addition to ProteomeXchange dataset (PXD) submissions for Arabidopsis, ProteomeXchange contains nearly one hundred PXDs for maize, providing an excellent opportunity to build a community proteomics resource also for maize. The current report describes the generation of the first Maize PeptideAtlas based on available maize MS/MS proteomics datasets in ProteomeXchange (cutoff date June 16 2023). In this report, we will first provide an analysis of what questions the maize community has addressed in these submitted MS-based proteome experiments and what genotypes and methodologies were pursued. Overall statistics on identified peptides, proteins and MS/MS match rates against the most recent B73 genome annotation by the maize community (https://www.maizegdb.org/genome/assembly/Zm-B73-REFERENCE-NAM-5.0) as well as previous annotations provides insight about proteome coverage and allows for comparison of B73 genome annotations (protein coding genes) across the different B73 versions in MaizeGDB and NCBI RefSeq. Finally, maize is an ancient polyploid with two subgenomes; a number of proteins are still represented by a gene on both subgenomes ^32–34^. This new Maize PeptideAtlas could be used to determine to what extent both duplicates are detected at the proteome level and if there is any subgenome bias.

This freely available Maize PeptideAtlas provides the global community with high quality, fully reprocessed MS-based proteome information together with its metadata. PeptideAtlas differs from other databases such as PPDB and Plant PTM Viewer in that the raw MS data from laboratories around the world are reprocessed. All identified peptides, PTMs, and MS/MS (tandem MS) spectra in PeptideAtlas are linked to the metadata collected from the PXDs, publications, and additional information from the submitting labs. PeptideAtlas is integrated with the maize community resource MaizeGDB ^5^.

## METHODS

### Selection and downloads of ProteomeXchange submissions

We downloaded all available raw files with information referring to maize or corn (*Zea mays*) from ProteomeXchange repositories with a cutoff date of June 16, 2023. Upon inspection, some of the PXDs were not further processed because i) they only contained recombinant maize proteins, ii) were mis-annotated as containing maize samples, iii) the data were only peptide mass printing (PMF) acquired on a MALDI-TOF instrument, or iv) raw files were missing. Supplemental Data Set S1 (and the abbreviated version Table 1) provides the list of all selected and processed PXDs, number of raw files and MS/MS spectra (searched and matched), genotypes of the samples, identified proteins and peptides, submitting lab and associated publication, as well as several informative key words.

**Table 1.** Summarizing information of the 74 selected PXD datasets for the maize PeptideAtlas build. This includes PXD #, publication, number of matched MS/MS spectra and % match rate, the number of identified proteins (canonical and groups of proteins), the number of matched distinct MS/MS peptides, the MS instrument, information about the sample (plant part, subcellular fraction, enrichment for PTMs, multiplexing by TMT or iTRAQ). An extended table with additional information is provided as Supplemental Data Set S1.

| PXD (until June 16, 2023) | Matched # MS/MS spectra | Matched MS/MS spectra (%) | # Distinct peptides at FDR 0.002 | plant part | B73, W22 or other genetic background | Instrument | Keywords |
| --- | --- | --- | --- | --- | --- | --- | --- |
| PXD001125 | 197,429 | 61 | 36,079 | root | A3237 & A3239 | Q Exactive | Waterlogging stress |
| PXD002027 | 1,446,304 | 28 | 225,325 | root - PM | FR697 | LTQ Orbitrap XL | plasma membrane, spectral counting, water stress |
| PXD002379 | 651,578 | 77 | 180,601 | endosperm | W64A and mutants | Q Exactive | opaque, non-zein, ER stress, UPR, lysine-rich |
| PXD002575 | 392,216 | 26 | 143,076 | root - PM | B73 | LTQ Orbitrap Velos | peri-arbuscular membrane, fungus |
| PXD002586 | 1306 | 9 | 274 | leaf | B73 | Q Exactive | Phosphoproteomic, Salicylic acid (SA). iTRAQ |
| PXD002853 | 72,480,339 | 22 | 2,827,041 | anther, cortex, ear, elongation zone, embryo, endosperm, germ kernel, internode, juvenile leaf, leaf, maturation zone, pericarp aleurone, pollen, pollen tube, pollen W23, primary root, secondary root, silk, spikelet, stele, tassel | B73; W23 | LTQ Velos | transcriptome, gene regulatory network, developmental atlas |
| PXD002877 | 728,095 | 17 | 65,968 | embryo | B73 | LTQ Velos | gene regulatory network, phosphoproteomics, RNA-seq, co-expression network, |
| PXD003634 | 165,189 | 53 | 30,560 | leaf | no info | Q Exactive | phosphoproteomics, drought |
| PXD006751 | 884 | 1 | 226 | leaf apoplast | Early Golden Bantam | LTQ | salicylic acid, peptide signaling |
| PXD007612 | 61,160 | 48 | 7,070 | etiolated coleoptiles | no info | TripleTOF 5600 | Golgi, flotation centrifugation, Free-Flow Electrophoresis |
| PXD007711 | 115,043 | 26 | 51,046 | root | Jing724 or D9H | Orbitrap Fusion | salt tolerance, comparative proteomic analysis, iTRAQ |
| PXD007802 | 241,531 | 27 | 63,394 | root | Jing724 or D9H | Orbitrap Fusion | salt tolerance, comparative proteomic analysis, iTRAQ |
| PXD007880 | 112,341 | 29 | 12,810 | leaf | B73 | LTQ Orbitrap | De-etiolation, ubiquitylation, C4 photosynthetic enzymes, ribosome, |
| PXD008186 | 78,407 | 41 | 50,635 | leaf | B73 | Q Exactive | rough dwarf disease, Rice black streaked dwarf virus, differentially expressed proteins, TMT |
| PXD008315 | 108,946 | 23 | 43,427 | seed | Honey Bantam | Orbitrap Fusion ETD | smoke |
| PXD008904 | 296,658 | 54 | 39,991 | leaf | P178 & Lx9801 | Q Exactive | Southern corn rust |
| PXD009627 | 3,523,743 | 62 | 358,100 | leaf | W22 - mu mutants | Q Exactive | Autophagy, proteome remodeling |
| PXD010582 | 11,304 | 44 | 4,971 | apoplastic fluid | Early Golden Bantam (EGB) | maXis QTOF | F24 PLCPs |
| PXD011039 | 53,349 | 27 | 18,246 | microsomes, tonoplast, vacuole | W22 | Q Exactive | microautophagy, aleurone, ER, receptors, TMT |
| PXD011363 | 7,469 | 9 | 1,800 | seed | DKC 2675 | Orbitrap Velos-Pro mass spectrometer | transformation; Genetically modified (GM) |
| PXD012022 | 86,856 | 31 | 15,243 | seedling | B73 | TripleTOF 5600 | Virus infection, MCMV, SCMV |
| PXD012344 | 127,356 | 34 | 50,291 | root | Chang 7-2 & TS41 | TripleTOF 5600 | Root, drought stress, varieties, iTRAQ |
| PXD012865 | 402,329 | 45 | 56,733 | leaf | no info | Orbitrap Fusion | protein complexes, SEC |
| PXD012897 | 515,891 | 40 | 120,323 | leaf | B73 | LTQ Orbitrap Velos; Q Exactive | De-etiolation, Phosphoproteome |
| PXD013124 | 97,429 | 17 | 33,959 | root apoplast | Early Golden Bantam (EGB) | LTQ Orbitrap Elite | papain-like cysteine proteases, activity-based protein profiling, root apoplast |
| PXD013606 | 384,210 | 44 | 140,145 | unknown | no info | LTQ Orbitrap | Phylogeny |
| PXD013891 | 90,473 | 12 | 10,438 | anther | A188 | Orbitrap Fusion ETD | Argonaute, Meiosis, Phosphorylation |
| PXD014407 | 3,936 | 17 | 2,475 | root | DKB 789 | LTQ Orbitrap | humic acids, label-free proteome, sustainable agriculture technology |
| PXD014409 | 107,728 | 31 | 42,290 | root | 8723, P138 | TripleTOF 5600 | salt, tolerance, iTRAQ |
| PXD014602 | 318,998 | 45 | 90,845 | root | B73 | LTQ Orbitrap Velos | Transcriptome, Alternative splicing, De-etiolation |
| PXD014603 | 6,288 | 7 | 2,927 | leaf | B73 | Q Exactive HF | Lysine-acetylation, De-etiolation, Photosynthesis |
| PXD015819 | 550,858 | 50 | 105,702 | leaf | hybrids ZD808 & ZD909 and their parents | Q Exactive | heterosis, hybrid |
| PXD016033 | 90,060 | 50 | 19,469 | anther mitochondria | Ces87-1Rf4Rf4 | Q Exactive | Cytoplasmic male sterility (CMS), restorer |
| PXD016743 | 105,085 | 37 | 47,620 | ear leaf | DengHai605 (DH605) | Q Exactive Plus | waterlogging, 6- benzyladenine, TMT |
| PXD016909 | 532,143 | 25 | 49,426 | root, shoot | Zheng58, Chang7-2 | Q Exactive | phosphoproteomics, salt stress |
| PXD017077 | 1,525,859 | 60 | 184,440 | leaf | W22 | Q Exactive | Autophagy |
| PXD017080 | 492 | 1 | 193 | leaf | B73 | Q Exactive | plant peptide |
| PXD017081 | 9,121 | 10 | 3,339 | leaf | B73 | Q Exactive | Non-canonical peptides, Peptidogenomics, Six-frame translation |
| PXD017781 | 10,716 | 20 | 4,378 | leaf, root | B73 | LTQ Orbitrap XL | biostimulants, protein hydrolase, TMT |
| PXD018939 | 50,682 | 12 | 15,864 | root | B73 | Q Exactive | insect, TMT |
| PXD019635 | 194,169 | 68 | 47,238 | ribosome | B73 | Q Exactive | 3'Ribo-seq, eQTL |
| PXD019715 | 48,898 | 10 | 4,235 | unknown | B73 | Q Exactive | quantitative peptidomics, endogenous peptides, discovery, diurnal variation |
| PXD019804 | 8,540,860 | 74 | 27,650 | leaf | hybrids of UH007 & 254 dent lines | Q Exactive | GWAS, drought |
| PXD021503 | 184,235 | 33 | 24,698 | seed | ES ZOOM YG | Orbitrap Fusion | grain protein quality, seed storage proteins |
| PXD022863 | 3,449,757 | 60 | 79,977 | seed | W22 - mu mutants | Q Exactive | autophagy, seed development, atg12 |
| PXD023426 | 624,150 | 46 | 124,220 | seedling | KR701 & hei8834 | Orbitrap Fusion | chilling, TMT |
| PXD024342 | 168,101 | 30 | 55,404 | stalk | B73 | Q Exactive Plus | stalk rot, F. graminearum |
| PXD024698 | 176,018 | 60 | 61,673 | silk | B73 | Q Exactive | Trichoderma, Induced systemic response, Fusarium |
| PXD025997 | 1,538,827 | 47 | 136,125 | seedling | B73 | Q Exactive | Small open reading frame (sORF), Small Peptide (sPeptide), Ribo-seq |
| PXD026096 | 264,215 | 48 | 61,838 | seedling | B73 (likely) | Q Exactive HF |  |
| PXD026269 | 97,356 | 25 | 9,727 | root | Anyu308 | Orbitrap Fusion Lumos | Label-free proteomics, Root water uptake, Aquaporins, Irrigation, Field capacity |
| PXD026369 | 485,163 | 50 | 35,319 | root | Sugar Buns | Q Exactive HF | metaproteomics, metaproteome, protein extraction, plant-microbe interactions, plant microbiota, microbiome |
| PXD026378 | 2,617,402 | 64 | 109,088 | leaf (3rd & 4th) | B73, MO17, RIL103, RILP39 | Q Exactive | rare alleles, UTR, SNP |
| PXD026755 | 9,526 | 3 | 350 | kernel | Suyunuo5 | Orbitrap Fusion | waxy maize, kernel, drought |
| PXD026760 | 157,082 | 73 | 34,962 | root | Zhengdan 958 | Q Exactive | glucose, photosynthesis, carbohydrate metabolism, phenylalanine biosynthesis, |
| PXD026853 | 949,105 | 63 | 35,963 | seedling | W22 (mu alleles mms21-1,2) | Q Exactive | mms21, SUMOylation, SMC5, |
| PXD027378 | 7,040 | 15 | 1,883 | seedling | LH244 | Q Exactive | ZmRR1, ZmMAPK8, phospho-sites |
| PXD027417 | 13,111 | 22 | 6,524 | leaf | Qingnong-11 | Q Exactive HF-X | malonylome, photosynthesis, Calvin cycle |
| PXD028027 | 366,923 | 45 | 79,171 | leaf | XY335 & HN178 | Q Exactive | low nitrogen |
| PXD028508 | 193,116 | 45 | 25,336 | seedling | Shaandan609 (91227 x Chang7-2) & Shaandan902 (K22 x K12) | Q Exactive HF | drought tolerance, photosynthesis, iTRAQ |
| PXD028987 | 24,708 | 20 | 2,197 | root | Gelber Badischer Landmais | LTQ Orbitrap Elite | Antioxidant, aquaporin, electron transport system, hypoxia, plant growth regulators, redox homeostasis |
| PXD029993 | 38,995 | 35 | 9,505 | root | B73 | LTQ Orbitrap | PFC, PFAS, hydroponics, persistent organic pollutants, stress, TMT |
| PXD030131 | 4,336 | 28 | 2,996 | stem | B73 & HDA101 | Q Exactive Plus | Fusarium graminearum, lysine 2-hydroxyisobutyrylation |
| PXD030655 | 16,841 | 6 | 5,767 | seedling | B73 | Orbitrap Fusion | Phosphoproteome |
| PXD033386 | 83,480 | 37 | 8,130 | root | Gelber Badischer Landmais | Q Exactive HF | hypoxia, soluble proteins |
| PXD033817 | 427,243 | 36 | 49,777 | root | Allure® | Orbitrap Fusion ETD | drought, rhizobiome, roots |
| PXD034262 | 405757 | 34 | 128450 | seedling | B73 | Q Exactive HF | cold stress, phosphoproteome, photosynthesis, spliceosome, TMT |
| PXD036545 | 546,327 | 38 | 104,977 | leaf (3rd) | B73 | Q Exactive HF | C4, phosphoproteome, kinase-substrate network, light, temperature |
| PXD040256 | 16,430 | 16 | 3,373 | xylem | no info | LTQ Orbitrap XL | Acropetal translocation, xylem, polycyclic aromatic hydrocarbons |
| PXD042776 | 1,597,376 | 35 | 82,594 | leaf | 98 maize inbred lines (including B73) | Q Exactive | association panel inbreds, iTRAQ |
| PXD044303 | 586,453 | 32 | 17,640 | chloroplast membranes, chloroplast stroma | W22 | LTQ Orbitrap | leaf development; chloroplast biogenesis, Bundle sheath, mesophyll |
| PXD045274 | 57,229 | 56 | 12,047 | pollen | W22 | LTQ Orbitrap | pollen, soluble proteome, membrane proteome |
| PXD045331 | 546,495 | 29 | 41,603 | leaf proplastid, leaf plastid nucleoid | W22 & B73 | LTQ Orbitrap | nucleoids, proplastids, chloroplasts, leaf development |
| PXD046098 | 964,018 | 31 | 54,810 | leaf, bundle sheath | B73 & M016 | LTQ Orbitrap | Microscopy, developmental gradient, bundle sheet, C4 differentiation |

### Extraction and annotation of metadata

For each selected PXD, we obtained information associated with the submission, as well as the publication if available. This information was used to determine search parameters and provide meaningful tags that describe the samples in some detail. These tags are visible for the relevant proteins in the PeptideAtlas. If needed, we contacted the submitters for more information about the raw files. To facilitate the metadata assignments and association to specific raw files, we developed a metadata annotation system to provide detailed information to each matched spectrum for the users of the PeptideAtlas, and these metadata can be viewed in the Maize PeptideAtlas.

### Assembly of protein search space

We assembled a comprehensive protein search space (Table 2) comprising the predicted *Zea mays* protein sequences from https://www.maizegdb.org/ for the maize cultivar B73 (Zm-B73-REFERENCE-NAM-5.0 (aka B73 RefGen_v5 or MaizeGDB B73 v5), Zm-B73-REFERENCE-GRAMENE-4.0 (aka B73 RefGen_v4 or MaizeGDB B73 v4), B73 RefGen_v3 or MaizeGDB B73 v3, and maize cultivar W22 (Zm-W22-REFERENCE-NRGENE-2.0), ii) UniProtKB Zm ^35^, iii) NCBI RefSeq v4 and v5 (https://www.ncbi.nlm.nih.gov/refseq) ^36^, iv) the mitochondrial genome (B37N genotype - NC_007982.1 original paper ^37^), v) the plastid genome (genotype NC_001666.2 see German PhD thesis of Fritsche 1988 ^38^, and vi) 500 contaminant protein sequences (*e.g.* keratins, trypsin, BSA) and GFP, RFP and YFP protein sequences commonly used as reporters and affinity enrichments frequently observed in proteome samples. The footnotes in Table 2 provide the www links and information about the format of the protein identifier for each source.

**Table 2.** The assembly of maize protein sequences from different sources used as the protein search space, and the respective number of total, distinct, and unique sequences in each source, as well as the sequence-identical intersection among sources.

### The Trans-Proteomic Pipeline (TPP) data processing pipeline

For all selected datasets, the vendor-format raw files were downloaded from the hosting ProteomeXchange repository, converted to mzML files ^39^ using ThermoRawFileParser ^40^ for Thermo Fisher Scientific instruments or the msconvert tool from the ProteoWizard toolkit ^41^ for SCIEX wiff files, and then analyzed with the TPP ^29, 30^. The TPP analysis consisted of sequence database searching with either Comet ^42^ for LTQ-based fragmentation spectra or MSFragger ^43^ for higher resolution fragmentation spectra and post-search validation with several additional TPP tools as follows: PeptideProphet ^44^ was run to assign probabilities of being correct for each peptide-spectrum match (PSM) using semi-parametric modeling of the search engine expect scores with z-score accurate mass modeling of precursor m/z deltas. These probabilities were further refined via corroboration with other PSMs, such as multiple PSMs to the same peptide sequence but different peptidoforms or charge states, using the iProphet tool ^45^. For most datasets in which trypsin was used as the protease to cleave proteins into peptides, two parallel searches were performed, one with full tryptic specificity and one with semi-tryptic specificity (except for PXD002853, PXD002877, and PXD006751 from LTQ instruments with a total of ∼90 million MS/MS spectra, which were searched only tryptic due to computational resource constraints). The semi-tryptic searches were carried out by default with the following possible variable modifications (5 max per peptide): oxidation of Met, Trp (+15.9949), peptide N-terminal Gln to pyro-Glu (−17.0265), peptide N-terminal Glu to pyro-Glu (−18.0106), deamidation of Asn or Gln (+0.9840), protein N-terminal acetylation (+42.0106), and if peptides were specifically affinity enriched for phosphopeptides, also phosphorylation of Ser, Thr or Tyr (+79.9663). For the fully tryptic searches, we also added oxidation of His (+15.9949) and formylation of peptide N-termini, Ser, or Thr (+27.9949) - we deliberately restricted these mass modifications to only fully tryptic (rather than also allowing semi-tryptic) to reduce the search space and computational needs. Formylation is a very common chemical modification that occurs in extracted proteins/peptides during sample processing, whereas His oxidation is observed less frequently, but nevertheless at significant levels ^46, 47^. In both semi-tryptic and fully tryptic searches, fixed modifications for carbamidomethylation of Cys (+57.0215) were used if samples were treated with reductant and iodoacetamide. Isobaric tag modifications (TMT, iTRAQ) were applied as appropriate. Four missed cleavages were allowed (RP or KP do not count as a missed cleavage). Several datasets were generated with a combination of LysC and trypsin, but no other proteases were used. Some of the datasets (PXD006751, PXD017080, PXD017081) contain the analysis of extracted peptidomes in which no protease treatment was used and these datasets were searched with ‘no enzyme’. Several experiments required special additional mass modifications due to the sample preparation. We have tabulated the exact mass modifications used for each experiment in detail in Supplemental Data Set S2 via a set of single-character keys (ABCD…) that denote each mass modification pattern; each dataset has a combination of keys to provide the complete set of parameters.

### PeptideAtlas Assembly

In order to create the combined PeptideAtlas build of all experiments, all datasets were thresholded at a probability that yields an iProphet model-based FDR of 0.001 at the peptide level. The exact probability varies from experiment to experiment depending on how well the modeling can separate correct from incorrect. This probability threshold is typically greater than 0.99. As more and more experiments are combined, the total FDR increases unless the threshold is made more stringent ^48^. The final iProphet model-based peptide sequence level FDR across all experiments is 0.001, corresponding to a PSM-level false discovery rate (FDR) of 0.0001. Throughout the procedure, decoy identifications are retained and then used to compute final decoy-based FDRs. The decoy-based PSM-level FDR is 0.00008, FDR for identified distinct peptides is 0.0015, and the final protein-level FDR is below 0.006. Because of the tiered system, quality MS/MS spectra that are matched to a peptide are never lost, even if a single matched peptide by itself cannot confidently identify a protein.

### Protein identification confidence levels and classification

For the PeptideAtlas projects, proteins are identified at different confidence levels using standardized assignments to different confidence levels based on various attributes and relationships to other proteins using a relatively complex but precise ten-tier system developed over many years for the human proteome PeptideAtlas ^49^ and recently also applied to the Arabidopsis PeptideAtlas ^17^. Details are provided in Table 2 of ^17^. We also simplified this ten-tier system to a simpler four category system which is more accessible to non-experts, and use this to summarize most of our findings for the maize PeptideAtlas, similar as we did for Arabidopsis ^17, 28^. In the simpler four-category system, proteins that have no uniquely mapping peptides but do not qualify as canonical (same as Tier 1) are categorized as ‘uncertain’, corresponding to the sum of tiers 2-6. Proteins are categorized as ‘redundant’ if they have only shared peptides that can be assigned to other entries and thus these proteins are not needed to explain the observed peptide evidence (tiers 7-9). Finally, all other proteins that completely lack any peptides observed at our minimum PSM significance threshold are categorized as ‘not observed’ (tier 10). For all protein identifications and categorizations, all peptides must first meet the stringent PSM threshold already described above. For both systems, the highest confidence level category is the “canonical” category (Tier 1), which requires at least two uniquely-mapping non-nested (one not inside the other) peptides of at least nine amino acids in length with a total coverage of at least 18 amino acids, as required by the HPP guidelines ^50^.

### Handling of gene models and splice forms

The protein coding genes in maize Ref_gen-v5 are represented by gene models (transcript isoforms), which are identified by the term ‘P00n’ after an underscore at the end of Zm identifier (*e.g.* Zm00001eb370310_P001). Protein identifier formats for the other protein sources are all different and listed in the footnotes of Table 2. We refer to the translations of these gene models as protein isoforms. Most protein isoforms are very similar (differing only a few amino acid residues often at the N- or C-terminus) or even identical at the protein level. It is often hard to distinguish between different protein isoforms due to the incomplete sequence coverage inherent to most MS proteomics workflows. For the assignment of canonical proteins (at least two uniquely mapping peptides identified), we selected by default only one of the protein isoforms as the canonical protein; this was the ‘P001’ isoform. However, if other protein isoforms did have detected peptides that are unique from the canonical protein isoform (*e.g.* perhaps due to a different exon), then they can be given tier 1 or less confident tier status depending on the nature of the additional uniquely mapping peptides (length and numbers). If the other protein isoforms do not have any uniquely mapping peptides amongst all protein isoforms (for that gene), then they are classified as redundant (tiers 7-9 in the more complex system).

### Integration of PeptideAtlas results in other web-based resources

PeptideAtlas is accessible through its web interface at https://peptideatlas.org. Furthermore, direct links for each protein identifier (B73 v5) are provided between PeptideAtlas and PPDB (http://ppdb.tc.cornell.edu/) and all PeptideAtlas peptides are provided in the UniProtKB feature viewer (https://www.uniprot.org/) with links back to PeptideAtlas. Matched peptide entries in PeptideAtlas are visable in the B73 v5 maize annotated genome as the “Maize PeptideAtlas” track (under “Annotations”) in JBrowse at MaizeGDB ^5^.

## RESULTS AND DISCUSSION

### Features of maize ProteomeXchange submissions

We downloaded all available raw files from 99 PXDs referring to maize, corn or *Zea mays* from ProteomeXchange with a cutoff date of June 16, 2023 (Supplemental Data Set S1). Upon inspection, some of the PXDs were not further processed because i) they only contained recombinant maize proteins, ii) were misannotated as containing maize samples, iii) the data were only peptide mass printing (PMF) acquired on a MALDI-TOF instrument, or iv) raw files were missing or corrupt. We excluded PXD00943 because it contained only data independent acquisition (DIA) as these have no MS/MS scans directly associated with MS precursor ions; all other PXDs used data dependent acquisition (DDA). While DIA datasets often have fewer missed ions per run by avoiding the stochastic precursor selection problems of DDA, FDR control is more challenging and uncertain due to the multiplexing of fragment ions ^51^. A large ensemble of DDA runs, especially when complex peptide mixtures are pre-fractionated (*e.g*. SDS-PAGE or HPLC), are more likely to achieve high coverage with low FDR than DIA. Nonetheless, there are many efforts to improve the processing of DIA ^52^ and we are starting to develop a mechanism to integrate DIA data into the PeptideAtlas build process. Of the 99 PXDs, 74 PXDs passed our inspection criteria, their raw MS/MS data were searched, and results incorporated into the first maize PeptideAtlas build. Table 1 provides a summary of these PXDs, whereas Supplemental Data Set S1A provides more detailed information for these PXDs and Supplemental Data Set S1B provides information on the rejected PXDs not used in the build. The first maize PXDs in ProteomeXchange were released in December 2012; however, there are older submissions to PRIDE (https://www.ebi.ac.uk/pride/), one of the oldest partners in the ProteomeXchange consortium, but these were not transferred to ProteomeXchange for technical reasons. We resubmitted three of these older, published datasets from the van Wijk lab ^53–55^ to ProteomeXchange such that these could be included in the Maize PeptideAtlas. Figure 1A shows the timeline of public availability of the maize PXDs in ProteomeXchange, most of which were submitted through PRIDE (70%), followed by iProX (22%), MassIVE (7%) and jPOST (1%). For most of these PXDs (90%), the MS data were acquired using an Orbitrap type instrument from Thermo Fisher Scientific (Figure 1B). Initially these Orbitrap instruments were mostly the early generation of LTQ Orbitrap models (Velos/XL/Elite), followed by many PXDs using one of the different versions of the Q Exactive instrument, as well as lower number of PXDs with more recent Orbitrap models (Lumos, Fusion, Exploris). The remainder of the PXDs were acquired on a variety of other instruments, *i.e.* TripleTOF 5600 and MaXis QTOF, LTQ and LTQ Velos (Figure 1B).

**Figure 1.**
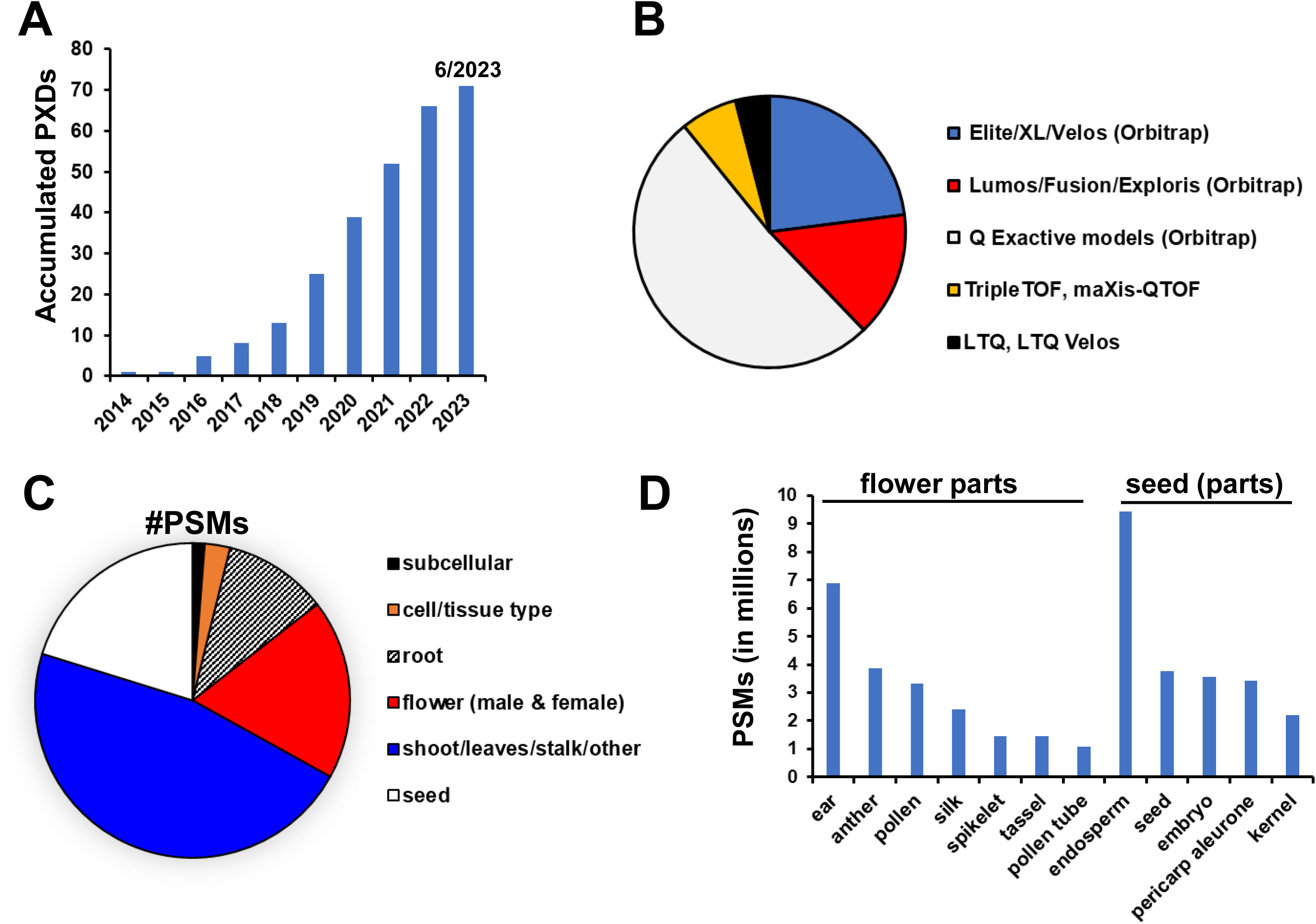
Features of maize PDXs used for the first maize PeptideAtlas build. (A) Cumulative PXD for maize in ProteomeXchange passing our selection criteria across the different years for maize. (B) Mass spectrometry instruments used to acquire data in the maize PXDs used in the maize PeptideAtlas grouped in different categories by vendor and model. (C) Distribution of PSMs of samples from different plant parts in the maize PeptideAtlas (D) Distribution of PSMs of samples for specific flower parts and seeds.

As indicated in Supplemental Data Set S1, submissions came from many different countries and continents, *i.e.* China (39 PXDs), USA (19 PXDs), Europe (12 PXDs), Japan (1 PXD) and South America (3 PXDs). The 74 PXDs address a wide range of topics, ranging from the response to various abiotic stresses (*e.g.* drought, waterlogging, salt) and biotic stresses (virus, bacteria, fungal infections), as well as biological phenomena such as de-etiolation, autophagy, development and differentiation (Table 1; Supplemental Data Set S1). Moreover, proteomes were extracted from a range of specific plant organs (*e.g.* root, leaf, pollen, silk, stem), or from specific subcellular locations and complexes (*e.g.* chloroplasts, apoplast, mitochondria, microsomes, plastid nucleoids) (Table 1; Figure 1C,D). 15 PXDs used multiplex labeling techniques (iTRAQ or TMT) for quantitative comparative proteomics (Table 1). Furthermore, 13 PXDs contained samples that were affinity-enriched to determine specific PTMs, including phosphorylation, ubiquitination, lysine acetylation, lysine 2-hydroxyisobutyrylation and lysine malonylation (Table 1). Furthermore, the PXDs cover a wide variety of maize cultivars and hybrids: i) 24 PXDs are exclusively from proteomes extracted from inbred line B73, ii) 7 PXDs are from proteins from the W22 inbred line, iii) many other PXD include samples from hybrids and their respective parents or commercial cultivars (*e.g.* Early Golden Bantam) (Table 1).

To build the PeptideAtlas, most PXDs were divided up in experimental sets of raw files in order to accommodate specific search parameters, treatments, cultivars, and sample types. This division of PXDs into experiments also allows the user to better explore the matched spectra, peptides and identified proteins within PeptideAtlas and its metadata annotation. A total of 213 experiments are assigned for these PXDs in PeptideAtlas, as described and summarized in Supplemental Data Set S2.

### Defining the protein search space

The maize community has made large investments in the B73 inbred line, including genome sequencing and annotations. Indeed, since the assembly and annotation of the first B73 genome ^2, 56^ there have been several successive B73 genome assemblies and annotations. The last three genome annotations by members of the maize community were RefGen_v3, RefGen_v4 (also named AGPv4) ^6^ and RefGen_v5 (Zm-B73-REFERENCE-NAM-5.0) ^7^ which each have their own set of (non-overlapping) gene identifiers (see Table 2 and footnotes for identifier formats). Furthermore, NCBI RefSeq also annotated these B73 physical assemblies. The informatics workflow for gene annotation of genome assemblies in RefSeq is different from those used for the maize community databases, and therefore result in only partially identical sets of protein sequences (Table 2).

In our search space we included the NCBI predicted maize proteomes GCA_000005005.6 (B73_RefGen_v4) with 131270 protein identifiers (including all isoforms) and the newer NCBI *Zea mays* Annotation Release 103 (based on Zm-B73-REFERENCE-NAM-5.0 GCF_902167145.1) with 72539 protein identifiers (including all isoforms; total 39756 protein coding genes) (Table 2). The search space also included the maize UniProtKB annotation with 63236 protein identifiers; this annotation is based on the annotation from EnsemblPlants and is continually updated through curation by UniProt. The inbred line W22 is popular in maize research, especially because of the availability of mutant collections with Mutator (Mu) and Dissociation (Ds) transposable element insertions for reverse and forward genetics studies ^8^. Indeed, eight PXDs used protein samples from this W22 background (Table 1) and we therefore also included the MaizeGDB W22 predicted protein sequences ^8^ in our search space (Table 2). In addition to the 10 nuclear chromosomes, maize (like all plants) has a small plastid (chloroplast) and mitochondrial genome, each with a much smaller set of protein coding genes ^37, 38^. Because several of the maize genome annotations do not include these organellar genomes, we included the predicted plastid- and mitochondrial-encoded protein sequences from NCBI RefSeq NC_001666.2 (plastid) and NC_007982.1 (mitochondria) and listed these as individual sources (Table 2; Supplemental Data Set S3 and S4). Finally, we also include a set of 500 protein sequences that represent common contaminants in proteome experiments, including keratins (from human skin and hair), trypsin (from autodigestion), proteins added to media such as BSA, various affinity tags, and fluorescent proteins (*e.g.* GFP, YFP, RFP and their variants).

Table 2 shows the number of total protein identifiers (including different isoforms or gene models) and unique protein sequences (unique here means that there was no other protein that was 100% identical, *i.e.* even a single amino acid difference between otherwise identical sequences would make them unique) for each genome annotation within our protein search space, as well as shared protein sequences between the different annotations. For instance, MaizeGDB B73 v5 has 72539 protein identifiers (including all splice forms/isoforms) of which 62559 are unique protein sequences within the v5 annotation. Just three protein sequences are only found in this source but not in any of the other sources, indicative of the overlap between sources. The total search space included 583838 protein identifiers (including all possible isoforms) representing 226062 unique maize amino acid sequences and the 500 unique contaminant sequences. In conclusion, we created a protein sequence search space that included the last three maize community annotations (MaizeGDB B73 v3-5), the last two NCBI-RefSeq maize annotations (NCBI RefSeq v4 and v5 *Zea mays* B73), UniProtKB *Zea mays* and the predicted maize plastid- and mitochondrial-encoded proteins (Table 2). This inclusive search space is a unique feature of PeptideAtlas (difficult to do by individual labs due the need for large computing resources) and will allow comparison of different B73 genome annotations and discovery of protein coding genes.

### Search results for the first Maize PeptideAtlas

Unless one uses *de novo* sequencing, MS/MS data can only lead to identification of peptides and proteins by searching these MS/MS data against an assembly of predicted, putative proteins. Proteins or peptides not represented in this protein search space cannot be identified. *De novo* sequencing is in principle possible and various software have been published (reviewed in ^57^), however it is hard to judge the quality of such searches and mapping back to proteins is challenging; searching different maize genome annotations is more efficient. Therefore, we assembled a comprehensive set of maize sequences from a variety of key sources as we described above (Table 2). Following downloading of PXD raw MS files, file conversions and sample annotations, the MS data were searched against this total protein space (see METHODS). We searched in several iterations to allow for the correct search parameters for each PXD and experiment (mostly variable and fixed PTMs). Especially PXDs that involved the use of tandem mass tag (TMT) or isobaric tags for relative and absolute quantitation (iTRAQ) labeling used for multiplexing and comparative proteomics (Table 1) required careful attention and verification of metadata.

The finalized searches and post-search processing for control of false discovery rates resulted in the matching of 120.4 million out of 444.8 million submitted MS/MS spectra, leading to the identification of 0.37 million distinct peptides (1.36 million different peptidoforms when counting different mass modifications). The overall match rate of MS/MS spectra to peptides (peptide spectral matches or PSMs) was 27%, but this match rate varied dramatically across PXDs from a few % to 80% (average and medium match rate is 34% and 33%) (Table 1). For those PXDs where we obtained a low match rate, we re-evaluated the search parameters to ensure that we did not overlook specific sample treatments that could affect the optimal search parameters (*e.g.* labeling techniques). The two PXDs with the lowest match rate were for peptidomics studies (leaf or apoplast) in which extracted proteins and peptides were not digested with trypsin or other enzymes (PXD006751 and PXD01780); this reflects the challenge to identify endogenous plant peptides.

To better understand the underlying data for this maize PeptideAtlas build, we calculated the frequency distributions of peptide charge state and peptide length for the PSMs (Figure 2A,B). The vast majority of matched spectra had a charge state of 2+ (63%), 3+ (29%) or 4+ (2%) and minor amounts of 1+ (5.6%) and 5+ (0.3%) (Figure 2A). We observed a wide range of matched peptide lengths, with seven amino acids being the shortest sequence allowed in the data analysis (Figure 2B). 99% of all matched peptides were between 7 and 35 aa in length with the most frequent peptide length of 15 aa. Figure 2C shows the number of identified distinct (non-redundant) peptides (irrespective of PTMs) as a function of peptide length (aa). The ratio between observed distinct peptides without missed cleavages and distinct peptides with missed cleavages and/or semi-tryptic peptides increased with peptide length (Figure 2C). This increasing ratio with peptide length illustrates that allowing for missed cleavages and semi-tryptic peptides increases average peptide length; it also improves protein sequence coverage and possible discovery of splice forms.

**Figure 2.**
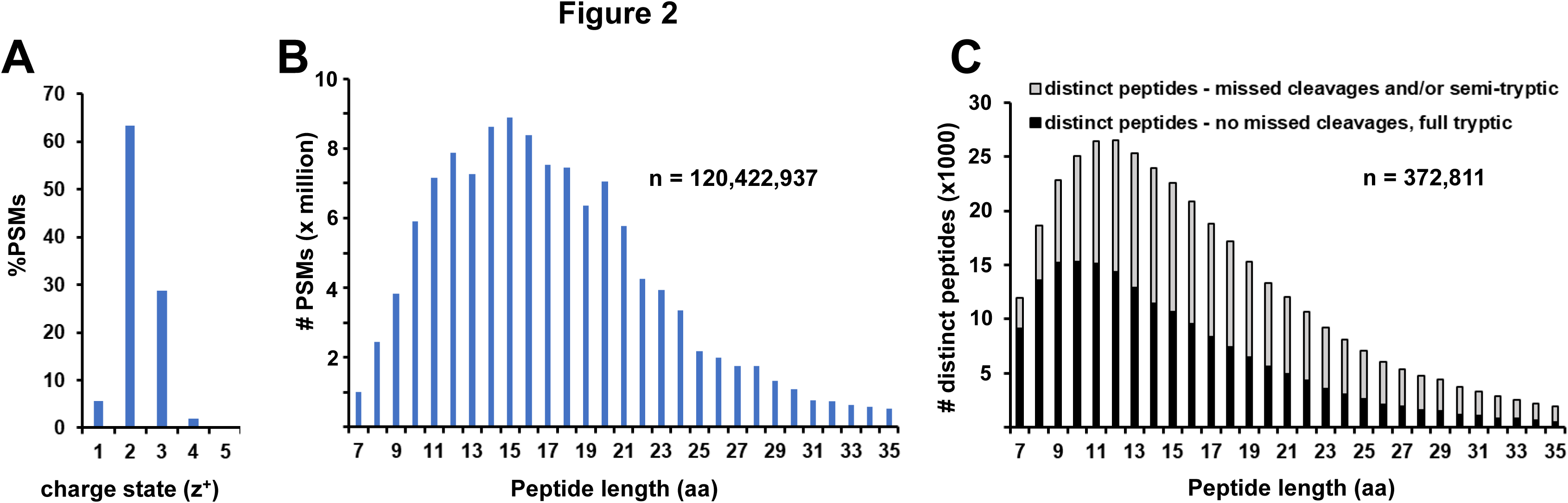
Key statistics of matched MS/MS data (PSMs) for this maize PeptideAtlas build. (A) Frequency distribution of PSMs for peptide charge state (z). (B) Frequency distribution of PSMs for peptide length (aa). Note that when R or K is followed by P, trypsin does not cleave and hence these are not counted towards missed cleavages. (C) Number of distinct peptides as a function of peptide length (aa).

Figure 3 shows the number of distinct (non-redundant) peptides (irrespective of PTMs) (Figure 3A) and distinct identified canonical (identified at the highest confidence level) proteins (Figure 3B) as function of the cumulative number of matched MS/MS spectra ordered by PXD identifier (from low to high or old to new) for this first Maize PeptideAtlas. Figure 3A shows that the cumulative number of distinct peptides rapidly increases with the first ∼2 million matched MS/MS spectra, followed by a gradual increase of cumulative peptides up to ∼90 million matched spectra. The matched spectra between 2 and 90 million (see arrow Figure 3A) all came from PXD002853 which sampled >20 tissue types using 10417 MS runs on a lower resolution LTQ Velos (Table 1, Supplemental Data Set S1) ^58^. The yield of additional new cumulative peptide per matched spectrum then rapidly increased because these came from PXDs of very diverse proteome sample sets and using higher resolution instruments. Figure 3B shows that after a rapid increase in newly identified canonical proteins based on the first ∼2 million MS/MS spectra, accumulation of additional newly identified canonical proteins gradually increased without showing any obvious saturation. Both Figure 3A and 3B indicate that future incorporation of newer PXDs into maize PeptideAtlas will likely lead to significant increase in newly identified distinct peptides and proteins. This contrasts the current status for the 2^nd^ release of the Arabidopsis PeptideAtlas https://peptideatlas.org/builds/arabidopsis/, where incorporation of additional and high quality PXD data set resulted in only small increases in distinct peptides and canonical proteins ^28^. Details about these maize PXDs, including how many new distinct peptides and proteins they identified, can be found in Table 1 and Supplemental Data Set S1.

**Figure 3.**
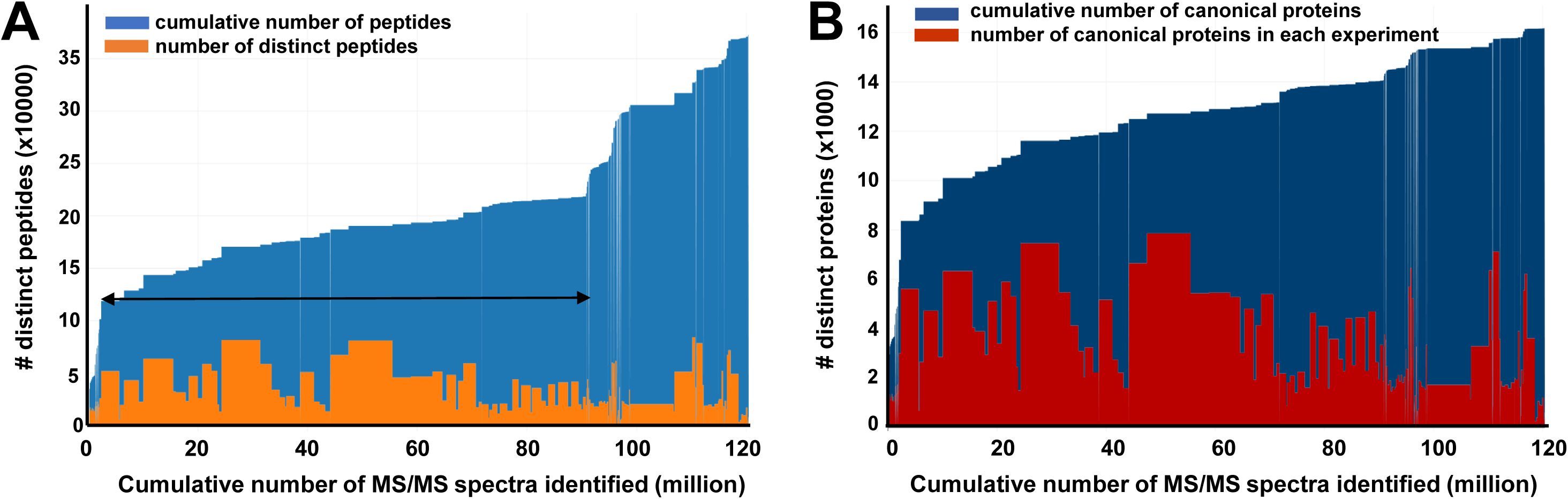
Number of distinct (non-redundant) peptides (left panel) and identified canonical proteins (right panel) as a function of the cumulative number of PSMs (peptide-spectrum matches) for the first maize PeptideAtlas. The cumulative count is ordered by PXD identifier (from low to high or old to new). The build is based on 211 experiments across the 74 selected PXDs, where each PXD may be decomposed into several experiments/samples when such information can be determined). The PSM FDR is 8.10-5. (A) Number of distinct (non-redundant) peptides as function of the cumulative number of MS/MS spectra matched. 372,811 distinct peptides are identified at a peptide-level FDR of 0.15%. Blue rectangles represent the cumulative number of distinct peptides as experiments are added to the build, whereas orange rectangles represent the number of distinct peptides in each experiment. The arrow from 2 million to 90.7 million MS/MS spectra are from PXD002853 acquired on a lower resolution LTQ-Velos instrument. (B) Number of distinct (non-redundant) canonical proteins as function of the cumulative number of MS/MS spectra matched at a canonical protein-level FDR of <0.006%. Blue rectangles represent the cumulative number of canonical proteins as experiments are added to the build, whereas red rectangles represent the number of canonical proteins in each experiment.

### Mapping peptides to the protein search space

Table 3 summarizes proteome coverage for the different maize protein sequence sources using redundant mapping of peptides. In such redundant mapping, peptides can match to more than one protein, which means that in many cases these proteins are not truly identified. Hence these initial numbers do not allow to make strong statements about which annotation version is better. 74% of all distinct protein sequences in B73 MaizeGDB v5 had peptides mapping to them (46134 out of 62559). Slightly lower percentages were observed for B73 MaizeGDB v3 and v4 (68% and 72%), but similar or somewhat higher for UniProtKB (74%), RefSeq v4 (78%) and RefSeq v5 (75%). 68% of all distinct protein sequences in W22 had matched peptides. We identified respectively 39 and 70 unique mitochondrial- and plastid-encoded proteins (Table 3) – these represent 26% and 76% of the predicted mitochondrial- and plastid-encoded proteins. This low percentage of identified organellar proteins is due to a vast over-assignment of protein coding genes in the RefSeq annotations (see below for discussion).

**Table 3.** Proteome coverage (redundant mapping) for the different protein sources in the search space.

| protein source | # protein coding genes | # protein ids including isoforms | # distinct protein sequences <sup>b</sup> | # observed proteins | observed distinct proteins (% of total) | # unobserved distinct proteins |
| --- | --- | --- | --- | --- | --- | --- |
| Core (B73 v5 _P001, mito, chloro) <sup>a</sup> | 40030 | 40030 | 38937 | 26019 | 67% | 12,918 |
| B73 v5 | 39756 | 72539 | 62559 | 46134 | 74% | 16,425 |
| mito RefSeq | 163* | 163 | 149 | 39 | 26% | 110 |
| chloro RefSeq | 111* | 111 | 92 | 70 | 76% | 22 |
| B73 v4 | 49376 | 143679 | 110536 | 79797 | 72% | 30,739 |
| B73 v3 | 38577 | 63241 | 57882 | 39368 | 68% | 18,514 |
| W22 | 40690 | 51716 | 42082 | 28439 | 68% | 13,643 |
| UniprotKB Zm | n/a | 63541 | 63232 | 46619 | 74% | 16,613 |
| RefSeq v5 | n/a | 57578 | 45919 | 34249 | 75% | 11,670 |
| RefSeq v4 | n/a | 131270 | 98578 | 77025 | 78% | 21,553 |
| CONTAM | n/a | 502 | 500 | 315 | 63% | 185 |

<sup>a</sup> Core proteome consists of one isoform per gene from the MaizeGDB proteome annotation from the Zea mays B73 version 5 genome assembly plus the mitochondrial and plastid proteomes from Refseq
<sup>b</sup> The number of distinct protein amino acid sequences (many different entries are sometimes sequence identical)
\* includes many pseudogenes

To better understand these peptide matches and to create a single Maize PeptideAtlas build, we assigned the matched MS/MS spectra in a hierarchical order with the predicted proteome of B73 MaizeGDB v5 having the first priority since we assume that the latest B73 genome annotation by the maize community is the most accurate version of the B73 genome annotation. Furthermore, the JBrowse tracks in maize GDB also show this latest version 5 projected onto the physical genome map, as well as annotated gene models with introns and exons. And as will be illustrated further below, all identified peptides in v5 are displayed in the JBowse track in MaizeGDB.

Many genes are represented by gene models (annotated as _P00n) due to possible alternative start and stop sites, different intron/exon boundaries. The result is that the number of predicted maize proteins far exceeds the number of predicted maize protein coding genes (Table 3). These gene models are represented by different protein identifiers (xxxx_P00n) but these protein identifiers might represent identical, protein sequences with just a few differences in amino acids or more substantially divergent protein sequences. Because it is often hard to obtain conclusive MS support for these highly similar protein isoforms (and of course impossible for identical protein isoforms), we created a ‘core proteome’ consisting of one protein isoform per protein-coding gene from MaizeGDB v5 plus the predicted plastid- and mitochondrial-encoded proteome. This predicted core proteome has 40030 different protein identifiers representing 38937 unique protein sequences. We observed peptides for 66.8% (26019) of these predicted proteins, *i.e.* for 66.8% of the protein coding genes we observed some MS-based evidence for their accumulation.

### Identification of the nuclear-encoded and organelle-encoded core proteome

Table 4 shows the proteins identified in the core proteome for each of the three confidence categories (canonical, uncertain, redundant – see METHODS for explanations) by nuclear chromosome (1-10), a small set of ‘scaffolds’ (*i.e*. predicted proteins that could not be assigned to a specific chromosome location) as well as the mitochondrial and plastid chromosomes. The identification rate (all three confidence levels) across the 10 nuclear chromosomes was very similar and ranged from 65% to 68% (Table 4). A lower percentage was identified for the unmapped scaffolds (45%) likely reflecting the lower quality or confidence of these predicted protein sequences. The number of core proteins identified at the highest confidence level (canonical; decoy-based FDR is <0.00006) was 16,178 which was 40% of the predicted proteome (Table 4).

**Table 4.**
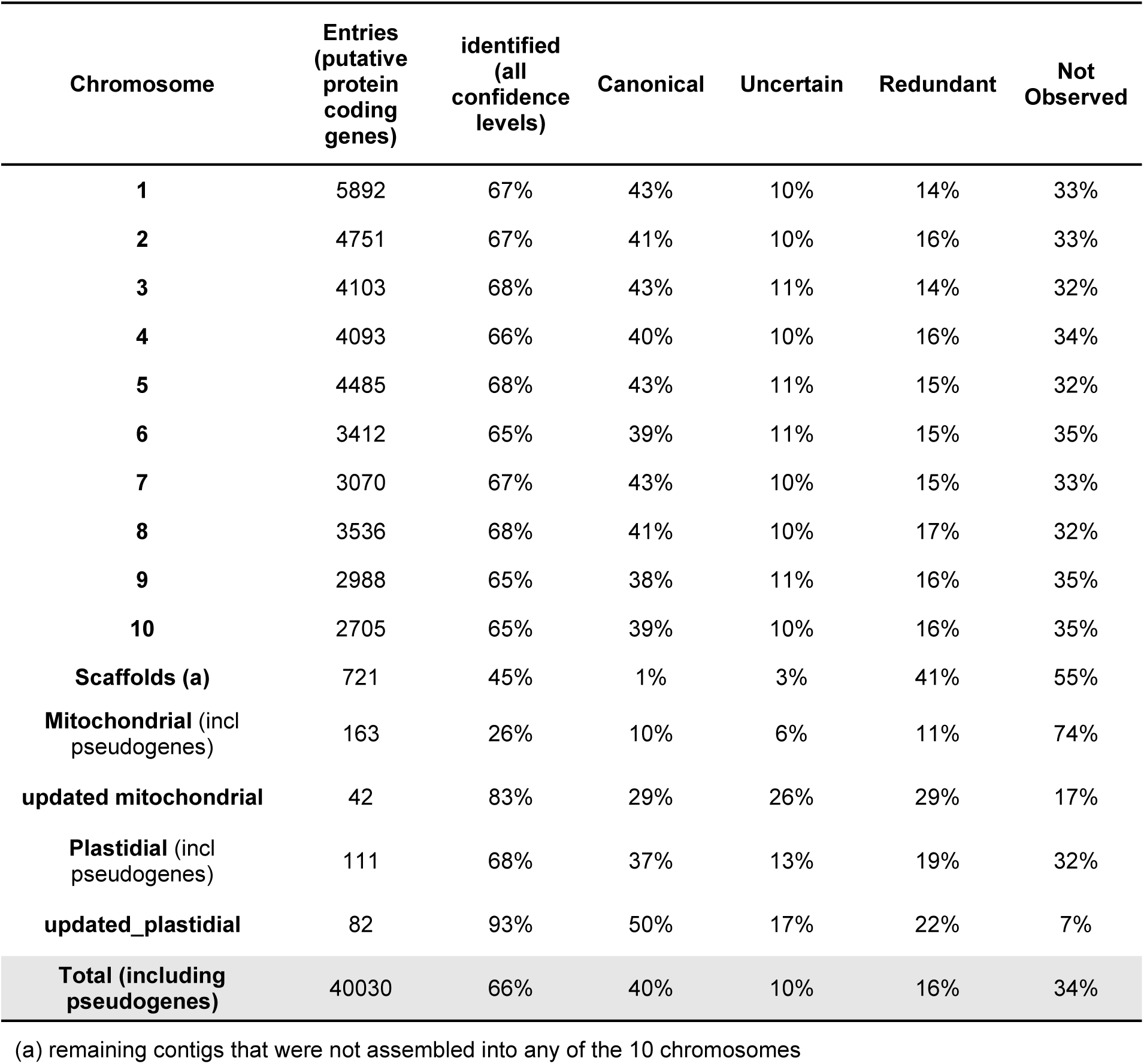
Identification status of the maize core proteome in the first maize PeptideAtlas build.

The predicted sets of plastid and mitochondrial encoded proteins are lacking in the MaizeGDB v5 since this annotation concerns only the 10 nuclear chromosomes (see Table 2). We therefore used the RefSeq annotations that are derived from ^37, 38^, which list 163 and 111 protein identifiers (one per gene) that encode for 159 and 92 unique sequences for mitochondrial and plastids, respectively. However, it is quite well established that the number of mitochondrial-encoded proteins in maize and higher plants typically includes around 33 conserved proteins (a dozen of which can be found duplicated) and the two optional ORFs T-urf13 and Orf355/Orf77 associated with cytoplasmic sterility ^37, 59^. We verified and updated the NCBI RefSeq annotations (see Supplemental Data Set S3) and list 42 annotated mitochondrial-encoded protein identifiers that include 5 pairs of identical sequences, and 3 pairs of closely related sequences. This also includes a pair of identifiers closely related to the protein with unknown function GRMZM5G892769_P01 in MaizeGDB v3. PSMs are reported for all but seven of these 42 proteins; the seven proteins are Nad4l, three of the cytc biogenesis proteins (Cmb, Ccmc, Ccmfc), Matr, Rpl6 and the ORF TatC/MttB. It is not surprising that no PSMs were obtained for these 7 proteins, since none of them have suitable predicted tryptic peptides that can be detected by MS (Supplemental Data Set S3). 121 of the protein identifiers are annotated as pseudogenes, based on the information from ^59^. Indeed, for 114 of these annotated pseudogenes, not a single MS/MS spectrum was found. For three pseudogenes, we found one small peptide, but this peptide was also matched to more confidently identified proteins. Interestingly, protein products for four annotated pseudogenes were identified at the canonical level (YP_588312.1, YP_588286.1, YP_588271.1, YP_588349.1) and closer inspection suggested that several of them appear *bona fide* identifications. These four proteins are glutharedoxin-like, two putative DNA polymerase B (PolB) and one putative RNA polymerase; for all four an identical protein was annotated in B73 v3 (See Supplemental Data Set S3). Previous reports stated that these polymerases genes are located on plasmids and that they are degenerate sequences ^59^. However, the data in PeptideAtlas indicate that these proteins clearly do accumulate; the significance of these proteins remains to be determined. Altogether, the information in PeptideAtlas for the mitochondrial-encoded proteins can be used to help generate a better predicted set of expressed mitochondrial-encoded proteins and could be combined with carefully assembled set of mRNA sequences *e.g.* by RNAseq and RiboSeq of purified mitochondria.

The 111 plastid identifiers include 29 pseudogenes (annotated as hypothetical proteins in RefSeq) and the remaining 82 identifiers representing established chloroplast proteins (Supplemental Data Set S4). Within these 82 identifiers are five sets of duplicated proteins because of duplication from the inverted repeats. PeptideAtlas identified all but six of these established proteins. These unidentified proteins are the small and hydrophobic integral thylakoid membrane proteins (between 29-40 aa in length and gravy index between 0.68 and 1.46) PetG, PetL, PetN of the cytb6f complex and PsbI and PsbJ of Photosystem II, as well as NdhG/Ndh6 (176 aa in length, five predicted transmembrane domains, 1.04 gravy index). These are likely not identified because they lack favorable peptides for detection by MS. The number of PSMs ranged from 0.6 million (RBCL) to just one (PsbM). Eleven plastid proteins had identical proteins in the MaizeGDB v5 annotation and two proteins (RbcL and ClpP1) had highly similar protein in the MaizeGDB v5 annotation, even if it is well-established that these plastid proteins are encoded by the plastid genome. These nuclear protein identifiers are likely pseudogenes from plastid genome fragments that are part of the nuclear genome. Finally, we also note that maize lacks plastid-encoded Ycf1, Ycf2 and accD observed in Arabidopsis plastids ^60^ and reported in the Arabidopsis PeptideAtlas ^61^.

### Identification of physiological PTMs in the core proteome

Plant proteins can undergo a number of post-translational modifications (PTMs) *in vivo* which can be irreversible (*e.g*. protein processing) or reversable (*e.g*. phosphorylation) (for reviews see ^62–65^). Many of these PTMs can be measured by mass spectrometry but often require specific affinity enrichment because they are present in sub-stoichiometric amounts or because they reduce the detectability of the peptide. A subset of the maize PXDs included such affinity enriched PTM datasets for phosphorylation (10 PXDs), ubiquitination (PXD007880), and three different lysine acylations, *i.e*. acetylation (PXD0146033), malonylation (PXD027417) and hydroxyisobutyrylation (PXD030131). The lysine ubiquitination and acetylation PTM analysis were both done in the context of the same course of maize leaf de-etiolation (B73) ^66, 67^. Supplemental Data Sets S5-S10 provide a summarizing information for the maize canonical core proteins for which the PeptideAtlas pipeline identified phosphorylation (S,T,Y) (Supplemental Data Set S5), N-terminal acetylation (Supplemental Data Set S6), lysine acetylation (Supplemental Data Set S7), lysine ubiquitination (Supplemental Data Set S8), hydroxyisobutyrylation (Supplemental Data Set S9), malonylation (Supplemental Data Set S10). PTM sites and number of PSMs per site within different p-value confidence intervals are also provided in these tables. The PTM viewer in PeptideAtlas allows for further investigation of all PTM-site assignments (including at lower confidence intervals) and associated MS/MS spectra as well as metadata.

For our analysis described here (Figure 4), we only considered those PTMs for which the site assignments were of high confidence, *i.e*. in the probability intervals 0.95 < p ≤ 0.99, 0.99 < p ≤ 1.00 and no choice (there was only one possible site for that PTM in the peptide). In total we observed 7419 canonical core proteins with one or more of these PTMs (46% of all 16178 observed canonical core proteins) and the overlap between proteins with phosphorylation, N-terminal acetylation or lysine PTMs (any of the four) is shown in the Venn diagrams (Figure 4A). This shows that the number of observed phosphoproteins was by far the highest at 6053, followed by 1845 proteins for which their N-terminus was acetylated (NTA), and 1484 proteins with one or more PTMs on the ε-amino of the lysine sidechains.

**Figure 4.**
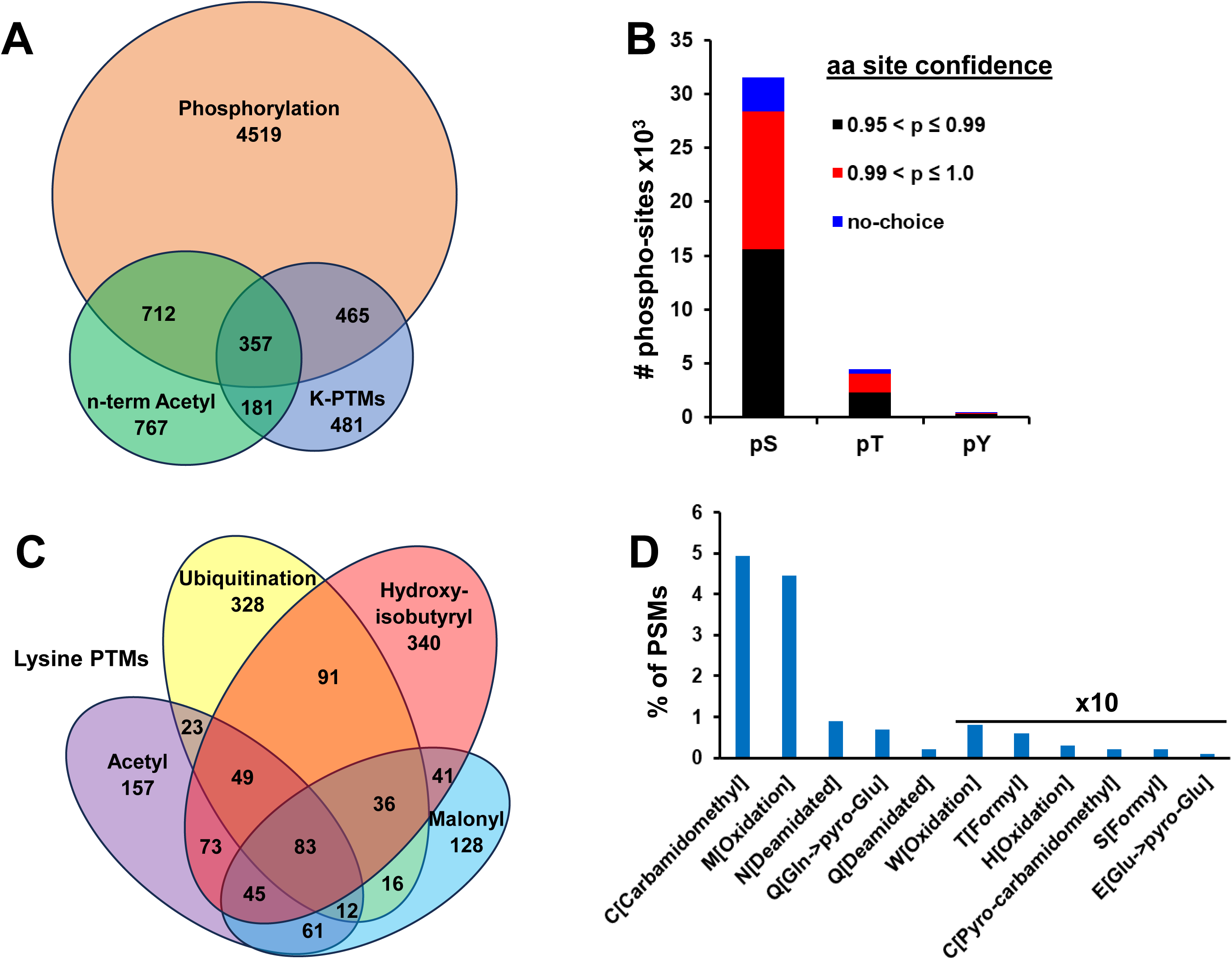
Canonical maize core proteins with one or more PTMs and frequency of PSMs with other mass modifications. (A) Overlap between proteins that carry one or more phosphorylations (S, T or Y), N-terminal protein acetylation (NTA), or one or more lysine modifications (K-ubiquitination, K-acetylation, K-malonylation, K-hydroxybutyrylation). In total there are 6053 proteins with one or more phosphorylations, 2017 proteins with N-terminal acetylation and 1484 proteins with one or more lysine side chain modifications (acylation or ubiquitination). (B) Number of serine, threonine and tryrosine phosphorylation sites (P-sites) at different p-value intervals for the core proteome. (C) Overlap between proteins with one or more lysine modifications. Colors: Acetylation - purple, Ubiquitination - yellow, Hydroxyisobutyrylation - red, Malonylation – blue. (D) Percentage of PSMs (of total) with mass modifications mostly due to sample preparations. Numbers are computed as the total number of PSMs that include at least one instance of the listed mass modification. Some PSMs contain more than one mass modification of the same type (not multiple counted) or different type (multiple counted).

Figure 4B illustrates shows the number of phosphosites at the three highest probability intervals for specific aa site assignment of the phosphate group for the canonical core proteome. The number of PSMs at the two highest probability intervals was about the same. There was total number of 36,451 phospho-sites (on average 6 p-sites per phosphoprotein) observed by 2.59 million PSMs. The ratio between pS, pT and pY sites was 86:13:1 which is consistent with other publications for meta-phosphoproteomics in Arabidopsis ^28, 68, 69^.

The lysine side-chain amine (ε-amine) can undergo a range of PTMs, including (poly)ubiquitination ^70^ and a dozen types of different acylations in particular in histones and a smaller number of types in non-histones ^71–73^. Malonylation, acetylation and 2-hydroxyisobutyrylation can be enzymatically catalyzed by lysine acyltransferases (KATs) and removed by lysine deacylases (KDACs). The functional significance for these acyl modifications is still very poorly understood. In contrast protein ubiquitination is typically a signal for degradation by the 26S proteasome, in particular in case of polyubiquitination and K48 linkages between the ubiquitin moieties. However, in most proteomics workflows, including in the PXD included here, only a di-glycyl (GG) remnants remains and hence information on whether the lysine was mono-ubiquitination or polyubiquitinated (and its linkage type) is lost. Figure 4B shows a Venn diagram for the proteins containing these different lysine modifications and ubiquitination. There was a significant overlap in proteins with more than one of these lysine PTMs, with 83 proteins have all four PTMs. It should be pointed out that we did not apply a minimum number of PSMs for any of these PTMs – hence closer inspection and consideration how the frequency of observation will be important if readers or users of the maize PeptideAtlas are interested to investigate specific proteins or PTMs. The Supplemental Tables S5-S10 and of course the Maize PeptideAtlas itself provide such information.

### Mass modifications typically due to sample preparation

In addition to biological PTMs (which require specific affinity enrichment for detection, except for N-terminal acetylation), the MS searches also include additional mass modifications that are induced during sample processing (see Methods). These modifications have generally very little biological relevance. The frequencies of these modifications can greatly vary between PXDs and experiments within PXDs depending on *e.g.* the use of organic solvents, urea, oxidizing conditions, temperature, alkylating reagent (alkylation of other residues than the intended cysteines), pH and use of SDS-PAGE gels. These mass modifications are included in the search parameters since many of these modified peptides would otherwise not be identified or lead to false assignments. The frequencies of these mass modifications (calculated as PSMs with the mass modification normalized to the total number of PSMs) are summarized in Figure 4D. This shows that the oxidation of methionine is by far the most frequent (4.5% of all PSMs), followed by deamidation of asparagine (0.9%), pyro-glutamate from N-terminal glutamate (0.7%) and deamidation of glutamine (0.2%), and very low levels (<0.1%) of tryptophan and histidine oxidation, formylation of threonine and serine, and threonine (0.7%), oxidation of proline (0.7%), pyro-glutamate from N-terminal glutamate and pyro-carbamidomethylation of cysteine. Carbamidomethylation of cysteine searched as fixed modification due to standard treatments with alkylating agents was 4.9%. These mass modifications are also visible in the PeptideAtlas web interface with viewable spectra and their interpretations.

### Identification of the non-core proteome

MS/MS spectra not matched to the core (maize GDB v5 – isoforms P001 + organelle-encoded proteins) were then assigned to subsequent sources, following the hierarchy as shown in Table 5. This table summarizes the hierarchical assignment of PSMs to the listed sources and identified peptides and proteins in different confidence categories. A listing and associated information for all non-core proteins for each of the sources identified at any of the confidence levels is provided in Supplemental Data Set S11A. Supplemental Data Set S11B provides a listing of all non-core proteins identified at the highest confidence level (canonical).

**Table 5.** Peptides assigned to proteins by hierarchy of sources ranging from core to DECOY (order from top to bottom), with each peptide is assigned only to the highest source possible and then not to any other source.

| Protein sources <sup>a,d</sup> | # Peptides | % Peptides | # PSMs | % PSMs | # Unique proteins (all confidence levels) | # Unique proteins (>=2 Distinct Peptides with >=3 PSMs) | Suppl Data Set S5 <sup>e</sup> |
| --- | --- | --- | --- | --- | --- | --- | --- |
| **Core (B73 v5_P001, mito, chloro)** | 338,111 | 89.8% | 113,683,979 | 93.5% | 23,302 | 17,096 |  |
| **B73 v5 (_P002 and higher)** | 16,550 | 4.4% | 4,150,816 | 3.4% | 4,547 | 2,186 | x |
| **UniProtKB Zm** | 629 | 0.2% | 308,998 | 0.3% | 121 | 65 | x |
| **B73 v4** | 8,360 | 2.2% | 1,494,636 | 1.2% | 2,445 | 912 | x |
| **B73 v3** | 1,017 | 0.3% | 232,734 | 0.2% | 454 | 101 | x |
| **W22** | 7,460 | 2.0% | 508,685 | 0.4% | 3,529 | 1,161 | x |
| **NCBI RefSeq (v4 & v5)** | 712 | 0.2% | 44,197 | 0.0% | 379 | 80 | x |
| **CONTAM <sup>b</sup>** | 3,280 | 0.9% | 1,086,243 | 0.9% | 266 | 153 |  |
| **DECOY <sup>c</sup>** | 528 | 0.1% | 37,173 | 0.0% | 522 | 4 |  |
| **Total** | 376,647 | 100% | 121,547,461 | 100% | 35,565 | 21,758 |  |

<sup>a</sup> Hierarchy refers to the order to which peptides are assigned to sources.
<sup>b</sup> Contaminants often found in samples, e.g. BSA, Keratin, trypsin, etc
<sup>c</sup> Decoys are all shuffled protein sequences in the search space; this enables accurate calculation of FDR.
<sup>d</sup> sources are defined in table 2
<sup>e</sup> Details for the non-core proteins identified at the highest confidence level (canonical) are provided in Suppl Data Set S5

As indicated in Table 5, matched peptides were first assigned to the core proteome. The vast majority of peptides (89.8%) and PSMs (93.5%) mapped to this core proteome showing that the protein space defined by the core proteome could explain the majority of matched MS/MS spectra across all samples from the PXDs selected for this first maize PeptideAtlas build. Most of the non-core matching peptides and the remaining PSMs, were assigned to the alternative isoforms (_P002 and higher) in maize GDB v5. These isoforms are generated by alternative splicing or alternative transcription or translation start sites, and provide an excellent resource to study the biological significance of these isoforms and quality of gene annotation. An example of identification of an alternative protein isoform for MaizeGDB v5 is demonstrated for Zm00001eb057170 encoding for a chloroplast-localized subunit of the thylakoid NDH complex (NDF5) involved in cyclic electron flow (Figure 5A). Sequence comparison of Zm00001eb057170_P001 (364 aa) and its second isoform Zm00001eb057170_P002 (411 aa) shows that the two isoforms are the same except for the C-terminal region, where they use different exons, and with isoform P002 in total 47 aa longer. There is no uniquely-mapping evidence in PeptideAtlas for the P001 isoform, but the P002 isoform is well supported by several uniquely-mapping peptides. This means that the default isoform P001 was likely not correct.

**Figure 5.**
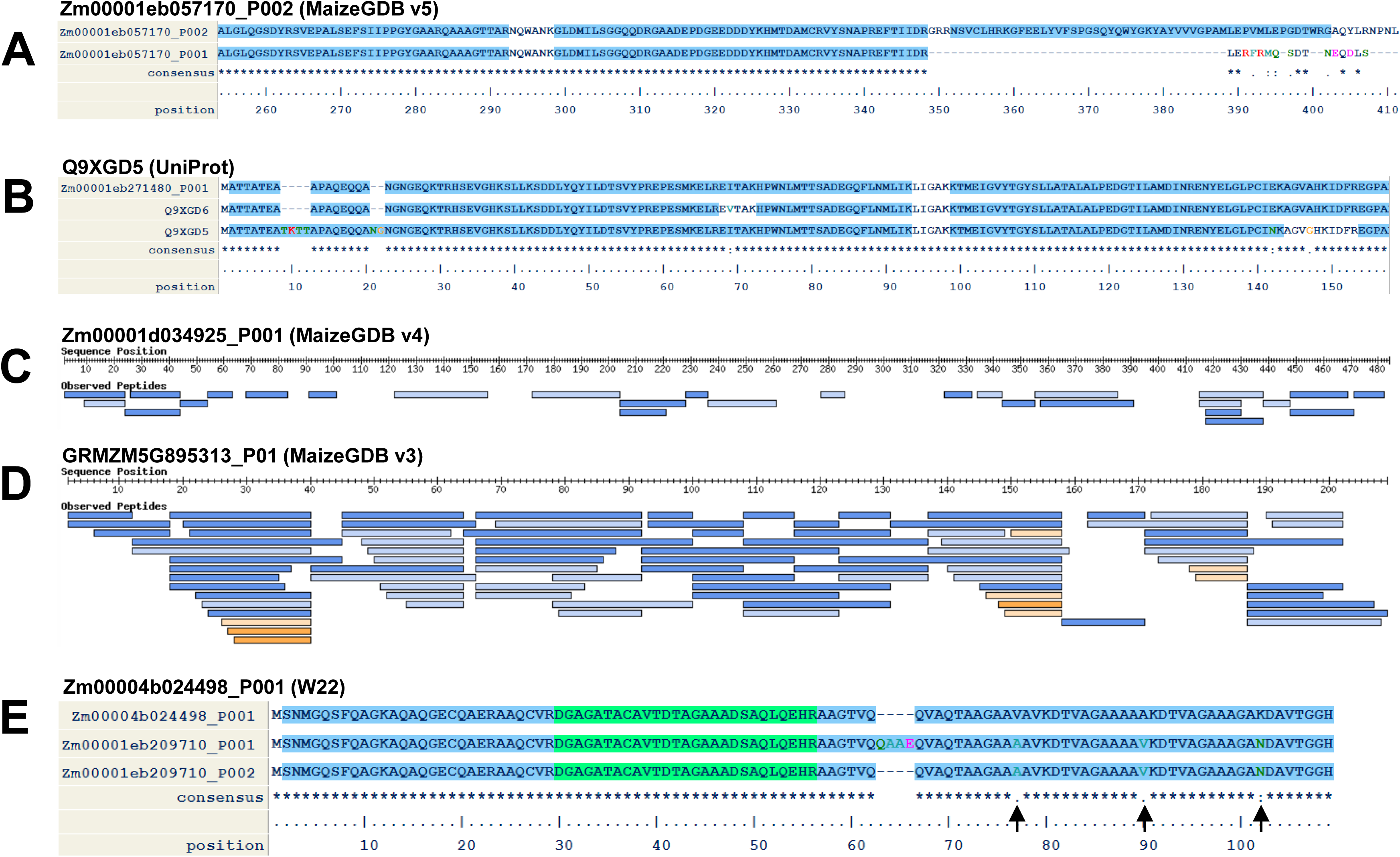
Examples of proteins that are well detected in the Maize PeptideAtlas that are not part of the MaizeGDB v5 core proteome. (A) Sequence comparison of Zm00001eb057170_P001 (NDH-dependent cyclic electron flow 5) and its second isoform Zm00001eb057170_P002 (both from MaizeGDB v5). The two isoforms are the same except for the C-terminal region, where they use different exons. There is no support in PeptideAtlas for the P001 isoform, but the P002 isoform is well supported by several peptides. (B) Sequence comparison of UniProt Q9XGD6 (Caffeoyl-CoA O-methyltransferase 1), UniProt Q9XGD6 (Caffeoyl-CoA O-methyltransferase 2) and Zm00001eb271480_P001, the closest entry in MaizeGDB v5. Although Q9XGD6 and Zm00001eb271480_P001 are highly similar (except for a V – I difference at position 69), there is no analog for Q9XGD6 in MaizeGDB v5. (C) Example of a protein (Zm00001d034925_P001) that is present in the MaizeGDB v4 proteome annotation but is absent from MaizeGDB v5. It is very well detected with 70% sequences coverage with 28 distinct uniquely mapping peptides, shown as blue rectangles. Darker shades indicate a larger number of PSMs. (D) Example of a protein (GRMZM5G895313_P01) that is present in the MaizeGDB v3 proteome annotation but is absent from MaizeGDB v4 and v5. It is very well detected with 100% sequence coverage (except for the initiating methionine) with 75 distinct uniquely mapping peptides, shown in blue (and a few short low-complexity peptides such as GGGGGGGR that also map to other proteins, shown in orange). (E) Example of a protein (Zm00004b024498_P001) in the MaizeGDB W22 proteome annotation that has many uniquely mapping peptides. It is very well detected with 8014 PSMs and 100% sequence coverage (except for the initiating methionine) with 36 distinct peptides, of which 24 are uniquely mapping to Zm00004b024498_P001. The skipped exon near position 65 is also encoded by a second isoform Zm00001eb209710_P002 from the B73 proteome. However, the single amino acid variants at positions 77, 90, and 102 are unique to the W22 sequence.

Only a small number of unique proteins (121) were identified in UniProtKB and these entries should provide explanations of the origin for these protein isoforms (Table 5). Identification of these 121 proteins only required 1 single peptide which likely results in a significant number of false identifications. When increasing the stringency to at least 2 matching peptides (these can be overlapping) and at least 3 PSMs, only 65 proteins were identified in UniProtKB (Table 5; Supplemental Data Set S11A). An example of a confidently uniquely identified protein from the UniProtKB annotation is Q9XGD6 (caffeoyl-CoA O-methyltransferase) (Figure 5B). The closest homology in MaizeGDB v5 is Zm00001eb271480_P001. Compared to this v5 homolog, Q9XGD6 shows four small regions with divergence, *i.e*. two small insertions (TKTT, AG) near the N-terminus confirmed with specific identified peptides and a point mutation (N instead of E) near the C-terminus which was also confirmed with a specific identified peptide.

Continuing down the hierarchical assignment, 8360 peptides (based on 1.5 million PSMs) were identified that mapped to MaizeGDB v4 but that did not map to v5 or UniProtKB. These peptides identified 2445 unique proteins (Table 5). Similar as described above for UniProtKB, increasing the stringency of identification to 2 peptides and at least 3 PSMs, only 912 proteins were identified in MaizeGDB v4 that were not already identified in v5 or UniProtKB (Tale 5; Supplemental Data Set S11A). Figure 5C shows an example of a protein (Zm00001d034925_P001; anthocyanin 5-aromatic acyltransferase) that is present in the MaizeGDB v4 proteome annotation but is absent from MaizeGDB v5. It is very well detected with 70% sequence coverage with 28 distinct uniquely mapping peptides, shown as blue rectangles. Darker shades indicate a larger number of PSMs. This shows that this protein isoform should be represented in a future B73 annotation.

454 proteins (not present in v5 or v4) (detected through 0.2 million PSMs) were identified in v3 across all confidence tiers and just 101 when requiring at least 2 matched peptides and at least 3 PSMs (Table 5; Supplemental Data Set S11A). Closer inspection of these unique proteins will help in the improvement of the v5 genome annotation. Figure 5D shows GRMZM5G895313_P01, an example of a protein (Glycine-rich protein 2b) that is present in the MaizeGDB v3 proteome annotation but is absent from MaizeGDB v4 and v5. It is very well detected with 100% sequence coverage (except for the initiating methionine) with 75 distinct uniquely mapping peptides, shown in blue (and a few short low-complexity peptides such as GGGGGGGR that also map to other proteins, shown in orange).

More than 3500 proteins (all confidence tiers) or 1161 proteins at the more stringent level that are not present in B73 v5, v4 or v3 were identified in the W22 annotation (Table 5; Supplemental Data Set S11A). Figure 5E shows that Zm00004b024498_P001 in the MaizeGDB W22 proteome annotation was identified with many uniquely mapping peptides. It is very well detected with 8014 PSMs and 100% sequence coverage (except for the initiating methionine) with 36 distinct peptides, of which 24 are uniquely mapping to Zm00004b024498_P001. The skipped exon near position 65 is also encoded by a second isoform Zm00001eb209710_P002 from the B73 proteome. However, the single amino acid variants at positions 77, 90, and 102 are unique to the W22 sequence.

What can we tell from these results about the quality of the three subsequent (v3, v4, v5) versions for the B73 annotations? The observation that the vast majority of matched spectra (96.9%) and matched peptides (89.9%) were observed for the protein space defined in v5 suggests that this latest genome annotation captures the majority of maize proteins. Version 3 and 4 together could only capture an additional ∼1.4% PSMs. To investigate these results further, we tested two other hierarchies of protein sources. Simply changing the order between v3 and v4 showed that v4 still captures more of the residual PSMs than v3 (0.8% for v4 versus 0.6% for v3; instead of 1.2% for v4 and 0.2% for v3). Starting the hierarchy with v3, then v4, followed by v5 core and v5 isoforms shows that v3 captured only 42.4% of the PSMs. But v3 & v4 together captured 76.0% and v5 still captured 22.5% of PSMs. Altogether, this shows that v5 is superior to v4, but a relatively small number (several hundred) of annotated proteins in v4 not present in v5, likely represent true protein forms. Closer inspection of the matched peptides in v4 and v5 will be needed to make decisions on the best annotation.

To help evaluate genome annotations of B73, the genome browser in MaizeGDB now includes a peptide track for v5. Similarly, peptide tracks for v3, v4 as well as W22 can be created to evaluate protein isoform annotations. We did annotate all experiments and PXDs for genotype (see Table 1) and in most cases the information for this meta-data annotation was available in the associated publications and/or PXD submission files. This showed that the majority of raw MS/MS data came from the B73 genotype, but many of the PXDs were other genotypes (Table 1). Therefore, when evaluating specific protein isoforms within or between genome annotations, one must be aware that the original genotype might have an impact on which isoform was identified. Because all matched MS/MS spectra are hyperlinked to the underlying meta-data it will in principle be possible to considering spectral quality, genotype and even sample treatment when evaluating protein annotations.

### How to use the maize PeptideAtlas build

This is the first effort to collect and reprocess all publicly available maize protein mass spectrometry data to provide a resource for the maize community. This resource will allow users to investigate proteins for their detection across many sample types, their protein sequence coverage which also might provide insight in post-translational processing (*e.g*. removal of the N-terminal signal peptide), accumulation of specific splice forms and possible PTMs. The maize PeptideAtlas home page interface (Figure 6A) displays the build summary and ‘clicking’ on the hyperlink “Full 2023-09 Build Summary” opens up a page with extensive information and query options, such as browsing by core proteins, canonical proteins or search by nuclear or organellar chromosome or other query options in the ‘Build Overview’ page (Figure 6B). There is a separate protein report page for each identified protein which includes: i) a protein map with observed peptides (and their significance for identification) and theoretical likely observable peptides, ii) a detailed PTM summary (*e.g.* phosphorylation, lysine or N-terminal acetylation), iii) listing with observed peptides with a range of properties, iv) list of experiments in which the protein was identified and hyperlinks to the experiment, PXD, publications. Hyperlinks to all matched spectra are available.

**Figure 6.**
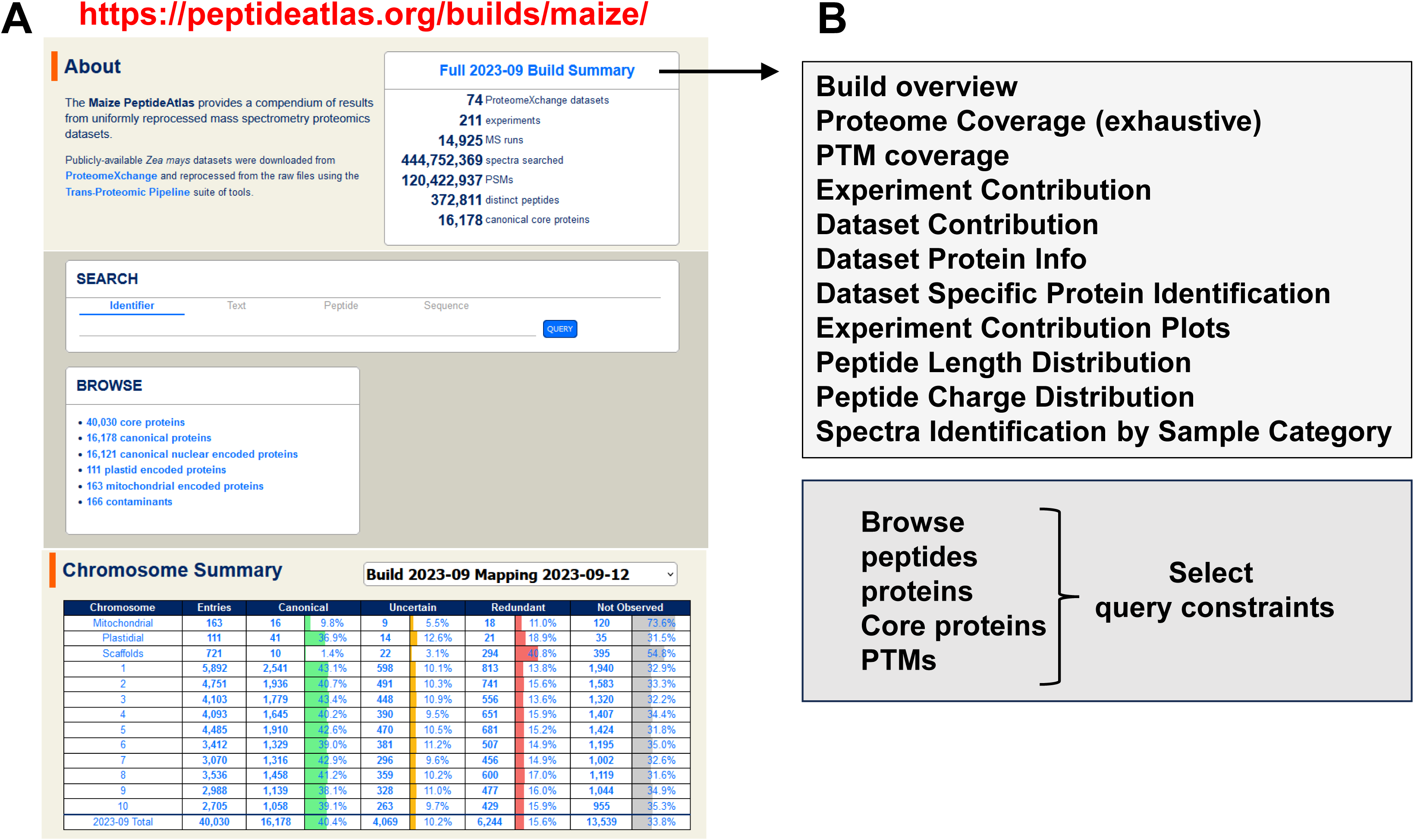
The maize PeptideAtlas home page (https://peptideatlas.org/builds/maize/) interface provides entry points to search and explore the database. (A) The home page display with the build summary, four different search functions, preloaded browse results, as well as a summary of identified and unidentified proteins for each of the 10 nuclear chromosomes, as well as plastid and mitochondrial chromosomes. (B) The hyperlink of Full 2023-09 Build Summary opens up a wide range of search options.

The maize PeptideAtlas home page interface (Figure 6A) also allows for four different search options: i) search by protein id, ii) search by free text, iii) search by amino acid sequence to find matched peptides (partial match is allowed), iv) search by amino acid sequence to map to the protein search space; the latter is like a FASTA file search. To illustrate these search functions (Figure 7), we use the maize homolog of a small chloroplast protein ClpS1, an N-recognin likely involved in the chloroplast N-degron protein degradation pathway ^74^. The protein identifier in maize GDBv5 is Zm00001eb041630 and the precursor is only 160 aa long (17 kDa). Searching by id (Figure 7A) resulted into one protein isoform (_P001) and provides a hyperlink to the specific protein report with information about possible and actual peptide sequence coverage, the detected PTMs, the observed peptides, in which experiments the protein was observed, and finally a mapping of all detected peptides across all experiments. Hyperlinks are available to explore the underlying data in far more detail. Using the text search (Figure 7B) with the input ‘CLPS1’ returns every maize protein where this text was found in the protein description – in this case Zm00001eb041630_P001 (Maize GDBv5), four ids from RefSeq v4 (ONM04608.1, ONM04607.1, AQK52459.1, AQK52458.1) and one from UniProtKB (B4FRF7). The results also show if the protein was identified and with how many observations (PSMs). CLPS1 in maize GDB v5 was identified at the canonical confidence level by 11 distinct peptides (with 1 to 98 PSMs; total 160 PSMs). Any of these peptides can be used to search the database. Using the example of peptide ‘AAPGKGGGVLDRPVEK’ (Figure 7C) returns this peptide, as well as two slightly longer peptides. This search query is useful if one identified a peptide in a proteome experiment and wants to explore to which proteins this matches or if PTM variants of this peptide have also been identified. Using that same peptide in the search function ‘Sequence’ (Figure 7D), returns any protein that contains that sequence, irrespective if that protein was identified. The maize PeptideAtlas home page interface (Figure 6A) also provides hyperlinks to browse for different groups of proteins, *e.g*. by chromosome, by identification confidence level (canonical, uncertain, redundant) or unobserved proteins.

**Figure 7.**
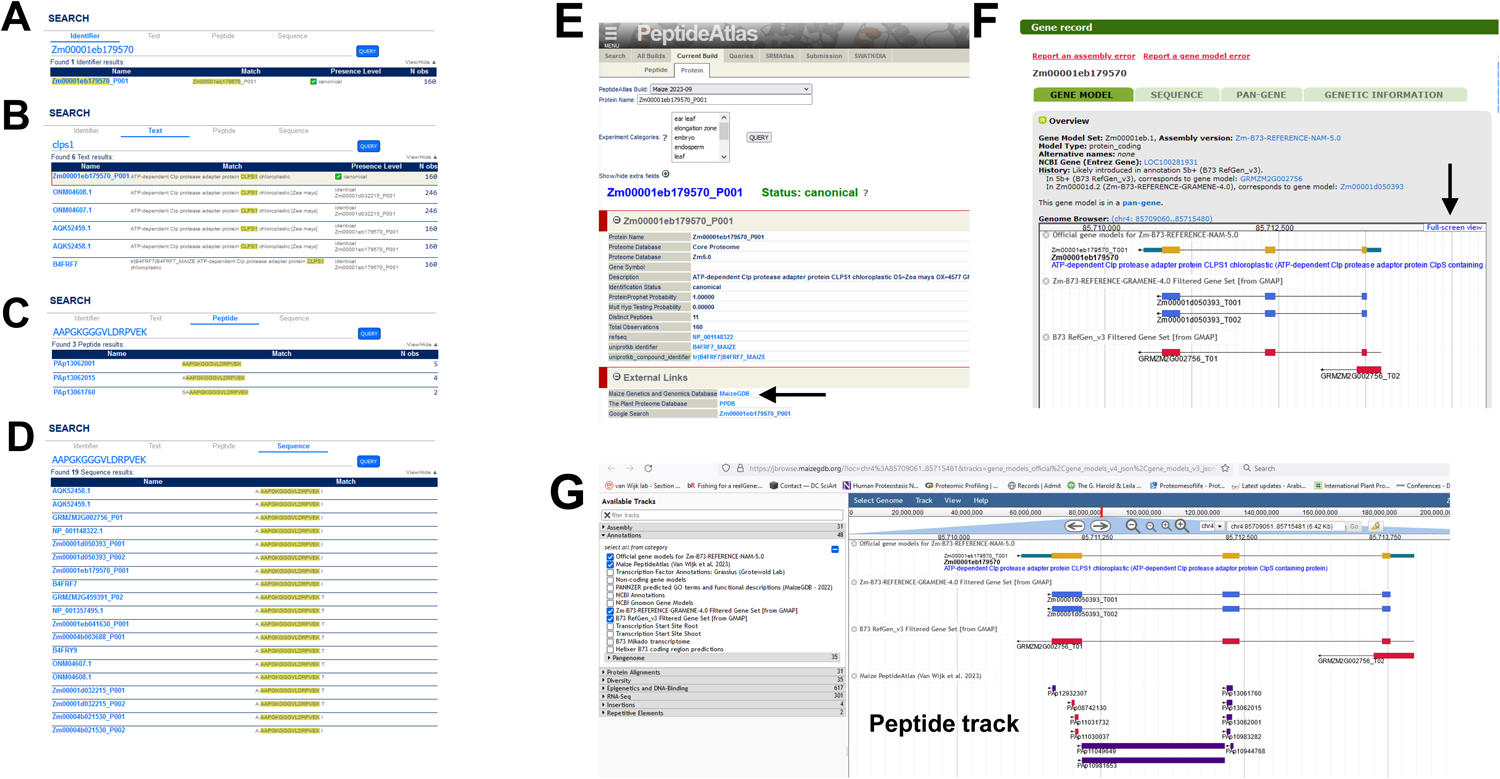
Search options in the maize PeptideAtlas interface and peptide display in the maize GDB genome browser. (A-D) Illustration of four different search options in PeptideAtlas as illustrated for the chloroplast protein CLPS1. (A) search by protein id, (B) search by free text, (C) search by amino acid sequence to find matched peptides, (D) search by amino acid sequence to map to the protein search space; this is similar as a FASTA search. (E) Partial view of the protein report page for CLPS1 in maize PeptideAtlas, including the link to maize GDB (see arrow). (F) Clicking on the hyperlink “MaizeGDB” (in panel E) opens the Gene record page in maizeGDB displaying the CLPS1 gene model in MaizeGDB v5, v4, v3. (G) Selecting the ‘Full screen view’ (panel F arrow) allows selecting additional tracks, include the peptide track with identified peptides from the maize PeptideAtlas.

A peptide track with matched peptides in the maize PeptideAtlas was added to JBrowse in MaizeGDB for reference genome v5 (Zm-B73-REFERENCE-NAM-5.0) and mapped peptides are hyperlinked back to PeptideAtlas such that information about the source of the peptide is easily available. Figure 7E-G illustrates this for CLPS1 Zm00001eb041630_P001. Figure 7E shows part of the protein report page in PeptideAtlas for CLPS1. Clicking on the hyperlink “MaizeGDB” (see arrow), opens the Gene record page in MaizeGDB (Figure 7F) – here displaying the CLPS1 gene model in MaizeGDB v5, v4, v3. Selecting the ‘Full screen view’ (Figure 7F see arrow) allows selecting additional tracks, include the peptide track with identified peptides from the maize PeptideAtlas (Figure 7G). In the case of CLPS1, this shows that the identified peptides match to the first two exons and also shows peptides that span both exons, supporting that the intron was indeed removed. The peptide track can thus help evaluate the quality of its predicted proteome, including alternative splice forms. Because different B73 genome assemblies and annotations were searched, the Maize PeptideAtlas can be used to evaluate the quality and relevance of these different genome assemblies.

### Comparison to the Arabidopsis PeptideAtlas

Compared to our most recent *Arabidopsis thaliana* PeptideAtlas (2023-10 release) ^28^ with an identification rate of 78.6% of the predicted proteome (counting one isoform per protein coding gene) across all confidence tiers, the percentage of identified maize proteins (counting one isoform per protein coding gene) in the latest B73 annotation was significantly lower (66.2%). This is likely a reflection of the lower amount of MS/MS data acquired on high resolution mass spectrometry instruments. The number of PXDs and associated publications for Arabidopsis is at least 5-fold higher than for maize and the latest Arabidopsis PeptideAtlas build includes 115 PXDs whereas this first maize build is based on 74 PXDs. Figure 3B shows that the cumulative number of detected proteins in the build continues to increase substantially even for the final dataset, whereas the analogous Figure 2B in ^28^ for the 2023-10 Arabidopsis build shows a clear saturation in the number of detected proteins. It is also quite likely that the higher complexity of the maize genome compared to Arabidopsis might further affect the identification rate in PeptideAtlas. As the number of high-quality maize PXD submissions will undoubtedly increase in the coming years, a future maize PeptideAtlas build will cover a higher percentage of the predicted proteome. However, the data contained in the current Maize PeptideAtlas can already be used to vastly improve the community annotation of the Maize reference proteome.

## Supporting information

Supplemental Data Set S1

Supplemental Data Set S2

Supplemental Data Set S3

Supplemental Data Set S4

Supplemental Data Set S5-10

Supplemental Data Set S11

## AUTHOR CONTRIBUTIONS

T.L. carried out all MS searches to create the PeptideAtlas build. Q.S. supported various steps in selecting and annotation of the maize sequence sources and established the links with the PPDB. Z.S. developed PeptideAtlas interface enhancements and assisted with the PeptideAtlas build process. L.M. developed PeptideAtlas interface enhancements and the dataset annotation tool. I.G., E.D., G.S., P.R. helped with incorporation of the metadata from the PXDs and associated publications into the PeptideAtlas internal annotation system. E.D. supervised the PeptideAtlas building process. K.J.V.W. contributed to the selection of PXDs and all aspects of specific plant biology-related issues. E.W.D. and K.J.V.W. developed this project, raised the funding, and wrote the paper.

## NOTES

The authors declare no competing financial interest.

## ACKNOWLEDGEMENT

This project was primarily supported by a grant from the National Science Foundation IOS #1922871 to K.J.V.W., E.W.D., and Q.S.. We thank the MaizeGDB team for their discussions and for incorporating the maize PeptideAtlas peptides into the maize B73 JBrowser.

